# A Multi-scale, Multiomic Atlas of Human Normal and Follicular Lymphoma Lymph Nodes

**DOI:** 10.1101/2022.06.03.494716

**Authors:** Andrea J. Radtke, Ekaterina Postovalova, Arina Varlamova, Alexander Bagaev, Maria Sorokina, Olga Kudryashova, Mark Meerson, Margarita Polyakova, Ilia Galkin, Viktor Svekolkin, Sergey Isaev, Grigory Perelman, Yaroslav Lozinsky, Ziv Yaniv, Bradley C. Lowekamp, Emily Speranza, Li Yao, Stefania Pittaluga, Arthur L. Shaffer, Danny Jonigk, James D. Phelan, Theresa Davies-Hill, Da Wei Huang, Pavel Ovcharov, Krystle Nomie, Ekaterina Nuzhdina, Nikita Kotlov, Ravshan Ataullakhanov, Nathan Fowler, Michael Kelly, Jagan Muppidi, Jeremy Davis, Jonathan M. Hernandez, Wyndham H. Wilson, Elaine S. Jaffe, Louis M. Staudt, Mark Roschewski, Ronald N. Germain

## Abstract

Reference atlases, molecular and spatial maps of mammalian tissues, are critical resources for discovery efforts and translational research. Their utility is dependent on operationalizing the resulting data by identifying cell types, histological patterns, and predictive biomarkers underlying health and disease. The human lymph node (LN) offers a compelling use case because of its importance in immunity, structural and cellular diversity, and neoplastic involvement. One hematological malignancy, follicular lymphoma (FL), evolves from developmentally blocked germinal center B cells residing in and trafficking through these tissues. To promote survival and immune escape, tumor B cells undergo significant genetic changes and extensively remodel the lymphoid microenvironment. Here, we present an integrated portrait of healthy and FL LNs using multiple genomic and advanced imaging technologies. By leveraging the strengths of each platform, we identified several tumor-specific features and microenvironmental patterns enriched in individuals who experience early relapse, the most high-risk of FL patients.

## INTRODUCTION

A chief aim of several international consortia is the construction of organ atlases from healthy and diseased human populations (Liu and Zhang, 2022; Regev et al., 2017; Rozenblatt-Rosen et al., 2020; Snyder et al., 2019). These efforts are focused on generating atlases from human organs and tissues using next generation sequencing (NGS) and spatial approaches such as multiplexed antibody-based imaging (Börner et al., 2021; Conde et al., 2022; Eraslan et al., 2022; Hickey et al., 2021; Jones et al., 2022; Suo et al., 2022). The lymph node (LN) offers unique challenges and opportunities due to its importance in host immune responses, structural and cellular diversity, distribution in >500 discrete units throughout the human body, and involvement in metastasis and hematological malignancies (Grant et al., 2020; Swerdlow et al., 2016).

Among the primary malignancies of the lymphatic system, follicular lymphoma (FL) is of special interest for multiomic examination. FL is a malignancy of germinal center (GC) B cell origin that extensively remodels the normal lymphoid tissue microenvironment. The disease trajectory of FL patients is remarkably heterogeneous, with many patients slowly progressing over several years, and a subset of patients experiencing an aggressive clinical course, often involving histologic transformation into diffuse large B cell lymphoma (DLBCL) or other high grade B cell lymphomas (Carbone et al., 2019; Casulo et al., 2015; Huet et al., 2018; Scott and Gascoyne, 2014). In rare instances, spontaneous remission may occur and is thought to require the generation of an antitumor response following immunologic stimuli, e.g., viral infection or injury (Potts et al., 2017). Lastly, patients who experience <u>p</u>rogression <u>o</u>f <u>d</u>isease within <u>24</u> months of initial treatment (POD24), termed early relapsers, are the subset with the shortest overall survival and should be considered the highest risk (Casulo et al., 2015; Casulo et al., 2021; Freeman et al., 2019; Maurer et al., 2016; Rodgers et al., 2021). Despite the urgent clinical need, a predictive tool and consensus treatment approach for early relapsers does not exist (Rodgers et al., 2021). Therefore, a greater understanding of the cell-intrinsic and -extrinsic factors governing progression and therapeutic outcomes is needed for risk-adapted management of FL patients.

FL B cells exhibit numerous genomic and epigenomic alterations that enable immune escape, apoptosis resistance, disease progression, and, in certain patients, histologic transformation (Carbone et al., 2019; Kridel et al., 2016; Lackraj et al., 2018; Scott and Gascoyne, 2014). The tumor microenvironment (TME) plays an integral role in supporting the survival of malignant cells throughout the course of disease. Genes expressed by dendritic cells (DCs) and macrophages have been associated with shorter patient survival, whereas genes expressed by T cells and macrophages are correlated with longer survival in FL (Dave et al., 2004). In a process described as “re-education”, FL tumor cells actively subvert the normal functions of non-malignant cells within the TME—T cells, follicular dendritic cells (FDCs), macrophages, DCs, and stromal cells—to support their survival and growth (Scott and Gascoyne, 2014). Furthermore, acellular elements and soluble factors warrant further investigation as they provide the scaffold and chemotactic signals required for cell migration within lymphoid microenvironments (Nagarsheth et al., 2017; Verdière et al., 2018).

Importantly, no single technical approach can sufficiently describe the cellular composition of normal LNs, nor identify transcriptomic and histological signatures associated with poor survival in FL patients. To overcome these challenges, we employed advanced sequencing and imaging technologies to generate both a molecular and spatial atlas of human LNs. Data integration yielded a unified description of lymphocyte populations, confirmed the extent of myeloid and stromal under-sampling from RNA sequencing (RNA-seq), and identified correlations between cytokine signatures and cellular communities obtained by the Iterative Bleaching Extends multipleXity (IBEX) imaging method (Radtke et al., 2022; Radtke et al., 2020). We additionally identified several cell intrinsic and extrinsic factors enriched in high-risk FL patients including enhanced B cell receptor (BCR) signaling and expansion of fibroblasts and extracellular matrix (ECM) components within the TME. In summary, this multiscale analysis of normal and malignant human LNs constitutes a valuable resource for discovery and translational research efforts.

## RESULTS

### Building a cell and tissue-level atlas using diverse lymphoid tissue sources

To create atlases of normal and FL LNs, we used state-of-the-art sequencing and imaging technologies (Figure 1). The study examined excisional LN biopsies from an ongoing clinical trial (NCT03190928), mesenteric LNs deemed grossly normal by an expert hematopathologist (nLN1-2), and a LN with reactive changes in the form of pronounced follicular hyperplasia (rLN1). The latter was included to address inflammation-associated changes versus tumor-induced molecular and morphological alterations in the LN. The clinical cohort included 7 FL (FL1-7) patients identified as non-progressors at least 2 years from study entry (Non-P), progressors within 2 years, or early relapsers (*) (Figure 1A, Table S1). Due to their small size and other technical constraints, non-FL LNs were limited to single cell RNA-seq (scRNA-seq) and IBEX imaging (Figure 1B). Excisional biopsies from FL patients, taken prior to any therapy, were subject to bulk RNA-seq, scRNA-seq, IBEX imaging, and additionally prepared as formalin-fixed, paraffin embedded (FFPE) samples for routine diagnostic pathology and multiplexed immunofluorescence (MxIF) imaging (Figure 1B). IBEX imaging was performed with the nuclear marker Hoechst and 39 antibodies targeting diverse cell types in regions of interest averaging 4-12 mm^2^ (Figure 1C-D). This approach revealed unique histological patterns that could be examined in larger regions (∼37-115 mm^2^) from clinically relevant FFPE samples using key markers identified from the IBEX data (Figure 1D). Several analytical methods were integrated, resulting in the construction of a molecular and cellular atlas of normal and malignant LNs across scales and modalities (Figure 1E, Table S2).

**Figure 1.**
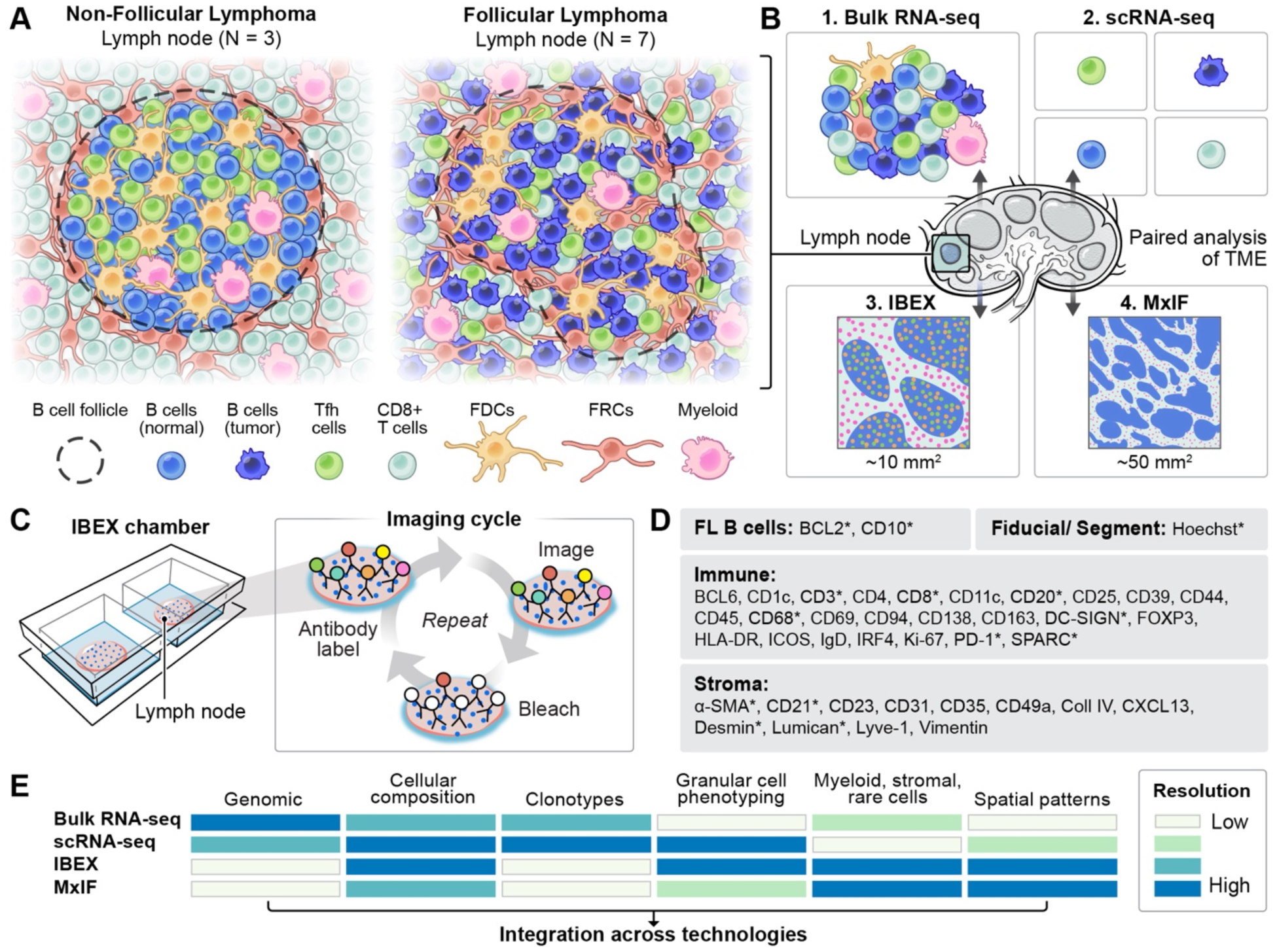
Construction of cellular atlases from human lymph nodes using multiple state-of-the-art omics and imaging technologies. **(A)** Tissue microenvironments (TME) were profiled in 3 non-follicular lymphoma (FL) samples and 7 excisional biopsies from FL patients. Follicular dendritic cells (FDCs), fibroblastic reticular cells (FRCs). Schematic shows enlarged and stylized secondary and neoplastic follicles. **(B)** Paired samples from normal and FL patients were profiled using multiple assays. Normal LNs were examined only by single cell technologies (scRNA-Seq and IBEX). Tissue microenvironment (TME), multiplex immunofluorescence (MxIF). (**C**) Schematic depicting IBEX imaging technique. **(D)** Protein biomarkers targeted with IBEX or MxIF (*) grouped by cell type. **(E)** Comparison of information provided by each technology. Resolution provided as an estimate only and based on analyses/results described in this work. For scRNA-seq: Clonotypes require TCR and BCR sequencing. Spatial patterns can be inferred.

### Genomic and transcriptomic characterization of FL heterogeneity

We first evaluated the genomic and transcriptomic landscapes of tumor B cells with whole exome sequencing (WES) and bulk RNA-seq. In addition to the expected translocations in the gene encoding the anti-apoptotic protein BCL2 (Table S1), diverse genetic lesions were identified in recurrently mutated genes previously associated with FL pathogenesis (Figure 2A) (Carbone et al., 2019). Alterations in chromatin-modifying genes such as *KMT2D, EZH2, ARID1A*, and *CREBBP,* as well as genes involved in cell migration and immune regulation *(CXCR4, TNFRSF14, CIITA)* (Carbone et al., 2019), were observed in progressors and non-progressors alike (Figure 2A). The cellular composition of FL samples was next evaluated by a recently developed deconvolution algorithm (Zaytcev et al., 2020) for data derived from bulk RNA-seq, a clinically feasible technology that provides expression data from diverse cell types (Figure 2B, Tables S3-S4). For all samples, B cells were the most abundant cell type, representing more than 60% of deconvolved cells from bulk analysis (Table S2). Myeloid and stromal cells, representing less than 2% of deconvolved cells per sample, were broadly classified as macrophages, monocytes, fibroblasts, and endothelial cells using lineage-specific genes (Tables S2-S4). In addition to analyzing the cellular composition of FL patient samples, malignant B cell receptor (BCR) sequences were identified from bulk RNA-seq based on the fraction of dominant immunoglobulin (Ig) sequences (Figure 2C) (Bolotin et al., 2017). Dominant clones were found in all patients except FL-1, a spontaneous remitter (Table S1). To further evaluate the clonal repertoire of FL B cells and overcome challenges arising from matching heavy and light chains from bulk suspensions, 5’ scRNA-seq was also performed (Figure 2C). The malignant Ig repertoires identified from bulk and scRNA-seq showed a similar monoclonal Ig distribution for all samples except for FL-1, confirming the utility of using bulk RNA-seq to deconstruct B cell clonotypes.

**Figure 2.**
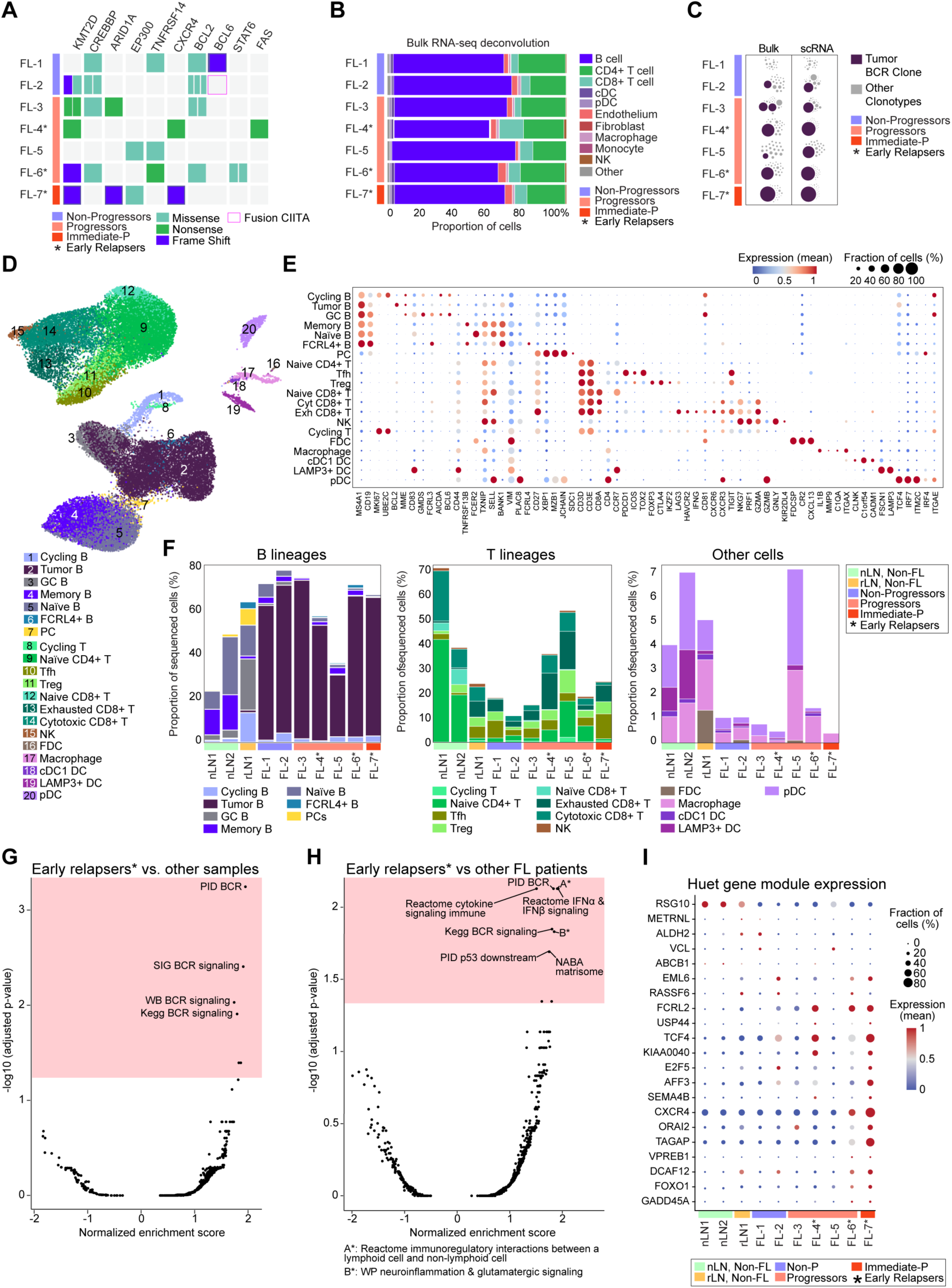
Cellular composition and gene expression patterns of normal and FL samples. **(A)** Genomic alteration landscape. Each line provides the detected mutations and fusions (cyan - missense mutation, green - nonsense mutation, violet - frameshift mutation, pink box - fusions) patients annotated based on progression status. **(B)** Cell composition reconstruction from bulk RNA-seq data using Kassandra algorithm. (**C**) B cell receptor calling from RNA-seq. Bubble corresponds to a unique or group of similar CDR3 sequences from heavy immunoglobulin genes. Size of circle corresponds to clonotype abundance. Bulk RNA-seq (bulk), scRNA-seq (scRNA) here and throughout. **(D)** Uniform manifold approximation and projection (UMAP) plot of 36,212 single cells from all samples. **(E)** Dot plot showing the expression of selected markers used for cell annotation of scRNA-seq clusters. Plasma cells (PCs), plasmacytoid dendritic cells (pDCs), Exhausted (Ex), and Cytotoxic (Cyt). **(F)** scRNA-seq frequencies of indicated cell types from each patient. **(G)** Gene set enrichment analysis of B cells from early relapsers (*) compared to all other samples plotted as enrichment score on the x-axis compared to the -log10 of the adjusted p-value on the y-axis. The pink box shows a cutoff of adjusted p-value < 0.05. Each point represents a gene set, with top scoring gene sets labeled. **(H)** Same as G but only comparing B cells from early relapsers to other FL samples. **(I)** Dot plot depicting dynamic expression of individual genes associated with Huet gene signature.

### Cellular composition of human LNs using scRNA-seq

To evaluate gene expression profiles at single cell resolution, we performed scRNA-seq on samples from FL patients. We additionally extended our studies to healthy LNs as a control and potential resource for LN atlas building efforts (Figure 2D-F, Figure S1A, Table S2). Distinct lymphoid populations varied in their relative abundance across samples (Figure 2F, Figure S1A, Table S2). These populations included naïve follicular B cells (*MS4A1, CD19, FCER2, SELL*), germinal center (GC) B cells (*CD83, GMDS, AICDA, BCL6, CD81),* cycling B cells (*MK167, UBE2C, CD81),* and tumor B cells *(BCL2, MME, TCF4)* found only in FL samples (Figure 2E-F). In-depth analysis of T cells revealed diverse subpopulations of CD4+ and CD8+ T cells including T regulatory cells (Tregs, *FOXP3, CTLA4, IKZF2, TIGIT*), T follicular helper cells (Tfh, *PDCD1 (PD-1), ICOS, TOX2, TIGIT*), and distinct populations of CD8+ T cells expressing several cytotoxic (*GZMA, PRF1, NKG7*) and exhaustion/activation-related (*LAG3, HAVCR2 (Tim-3), TIGIT*) markers. These ‘Exhausted CD8+ T cells’ were nearly absent from non-FL LNs but enriched in rLN1 and LNs from FL patients, suggesting chronic inflammatory reactions in these tissues (Figure 2F). Evaluation of myeloid and stromal populations by scRNA-seq frequently requires specialized protocols for tissue dissociation and cell enrichment, increasing the risk for altered gene expression profiles (Waise et al., 2019). To minimize these artifacts, we performed single cell analysis on suspensions obtained with limited intervention. Although representing a small fraction of total cells, pDCs (*IRF7, ITGAE, ITM2C, PLAC8, TCF4*), macrophages (*C1QA, IL1B, ITGAX*), cDC1 DCs (*C1orf54, CADM1, CLNK*), and FDCs (*CR2, CXCL13, FDCSP*) were profiled at varying frequencies across normal and FL samples (Figure 2D-F, Table S2).

### scRNA-seq derived gene signatures of FL patients

We next performed gene set enrichment analysis (GSEA) of scRNA-seq populations to identify the biological processes driving early progression and relapse in FL patients (Korotkevich et al., 2021). We first explored the gene expression patterns distinguishing B cells in early relapsers from B cells in all other samples. The top pathways upregulated in early relapsers all involved BCR signaling (Figure 2G). Several shared genes were identified as the leading-edge subset, defined as high scoring genes accounting for the enrichment signal (Korotkevich et al., 2021). These included mRNAs encoding for *CD19, BLNK, SYK, and LYN* (Table S2). We next compared the B cells from early relapsers to the B cells from all other FL patients. As before, the B cells from early relapsers upregulated components of BCR signaling pathways along with molecules involved in cytokine signaling, immune activation, and immunoregulation (Figure 2H, Table S2). We additionally observed enrichment of a gene set associated with glutamatergic signaling including transporters (*SLC38A1, SLC2A3, SLC38A2, SLC2A1*) and enzymes (*GLUL, GLS*) involved in glucose and glutamine metabolism (Table S2) (Wang et al., 2020). Quite surprisingly, the B cells of early relapsers exhibited high expression of genes involved in extracellular matrix (ECM) formation including those encoding growth factors, ADAMs, annexins, and galectins (Figure 2H, Table S2) (Naba et al., 2012).

In addition to the unbiased approach described above, we evaluated gene signatures correlated with poor outcome in FL patients (Huet et al., 2018) as well as IRF4-associated molecular signatures dysregulated in other hematological malignancies (Wang et al., 2014) (Figure 2I, Figure S1B-D). As expected, B cells from the early relapsers had elevated levels of transcripts from genes correlated with a high risk of progression (Figure S1B) (Huet et al., 2018). However, considerable interpatient heterogeneity was observed in the expression of individual genes summarized by the Huet module (Figure 2I). Within the IRF4-associated module, the early relapsers uniformly showed increased expression of *FOXP1* (adjusted p-value < 0.0001, log fold change = 1.07) (Figure S1C). This transcription factor has been shown to predict adverse failure-free survival (FFS) in FL patients treated with rituximab and chemotherapy (Mottok et al., 2018), the treatment regimen these patients received following the initial biopsies profiled here (Table S1). Of the three early relapsers, FL-4 and FL-7 displayed the highest aggregate expression of IRF4-associated genes as shown via violin plot (Figure S2D). These data demonstrate the power of scRNA-seq to identify unique patterns of gene expression in malignant cells from FL patients including those with poor prognoses and treatment failure.

### Assessment of clonotypes and *N*-glycosylation sites within variable regions of Ig genes

We next examined Ig clonotypes in our study using directed amplification and sequencing of BCRs. Retention of surface Ig expression is critical for malignant cells as it provides a mechanism of antigen recognition and survival signaling in the TME. While ∼40-50% of FL B cells are reported to undergo isotype switching to IgG, IgM is frequently observed in early stages of FL and thought to favor GC reentry (Küppers and Stevenson, 2018). The majority of progressors expressed IgM heavy chains except for FL-5, whose tumor cells had class switched to IgG (Figure S2A). We observed that all dominant FL BCR clonotypes have at least one, and in some cases several, *de novo N*-linked glycosylation sites in their heavy and/or light chain variable regions, regardless of progression status, as is typical of FL B cells (Zhu et al., 2002) (Figure S2B). Previously, the interaction between glycosylated BCRs on FL B cells and dendritic cell-specific intercellular adhesion molecule-3-grabbing non-integrin (DC-SIGN) has been shown to stimulate BCR survival signals in malignant cells (Amin et al., 2015; Coelho et al., 2010; Küppers and Stevenson, 2018). Given our observation of BCR signaling-related gene expression in early relapsers (Figure 2G-H), we believe this mechanism to be active in B cell clones from these patients. In summary, bulk and scRNA-seq revealed diverse cell-intrinsic factors governing survival and progression in FL patients.

### IBEX imaging provides a spatial atlas of the tissue microenvironment

To provide a spatial context for the cellular heterogeneity observed across normal and FL LNs, we performed 40-plex IBEX imaging with 39 markers specific for various cell lineages plus the nuclear label Hoechst (Figure 1D, Figure 3A). For accurate identification and quantification of cell types *in situ*, individual cells were segmented using several membrane markers and Hoechst (Figure S3). A deep learning-based approach was applied that assigned cell types based on the cosine similarity between cell marker expression and vectors from reference probability tables (Figure S3A-C, Table S5). Using this approach, 37 phenotype clusters were identified, resulting in a single cell proteomic dataset of 1.8×10^6^ cells from all 10 samples (Figure 3B, Table S2). Phenotype clusters were annotated into cell types based on protein biomarker expression, visualized using the data dimensionality reduction method Uniform Manifold Approximation and Projection (UMAP), and named according to their dominant marker expression and, whenever possible, relevant cell ontologies (Börner et al., 2021; Jorgensen et al., 2021) (Figure 3C). Our segmentation workflow produced accurate quantification of lymphocytes and, importantly, provided an independent confirmation of the rich diversity of T cell populations identified by scRNA-seq (Figure 3A-E). Furthermore, IBEX imaging revealed a very high level of heterogeneity among B cell populations across FL samples (Figure 3D, Figure S4A-B, Table S2). Although myeloid and stromal cell populations were identified using the segmentation approaches outlined here (Figure 3B-C), these cell types pose significant challenges due to their complex morphology (Germain et al., 2022; Risom et al., 2022). To overcome these challenges, we developed a method for profiling myeloid and stromal cells based on the creation of image masks that use pixel level data instead of discrete segmented cells (Figure S3D). Cell quantification of multiple myeloid and stromal lineages was performed using the indicated markers and normalized across samples by area imaged (Figure 3F-I, Table S2). Using this approach, we evaluated the location and relative abundance of blood endothelial cells, lymphatic endothelial cells (LECs), FDCs, and several fibroblast types including CD49a+ cytokine/chemokine-producing fibroblastic reticular cells (FRCs) recently identified in FL LNs (Mourcin et al., 2021) (Figure 3F-I, Figure S4C). Quantitative imaging revealed these cell types to be far more abundant than appreciated by methods employing routine tissue dissociation approaches (Figure 3G-I, Table S2). The resulting data underscore the importance of studying complex cell types in intact tissues.

**Figure 3.**
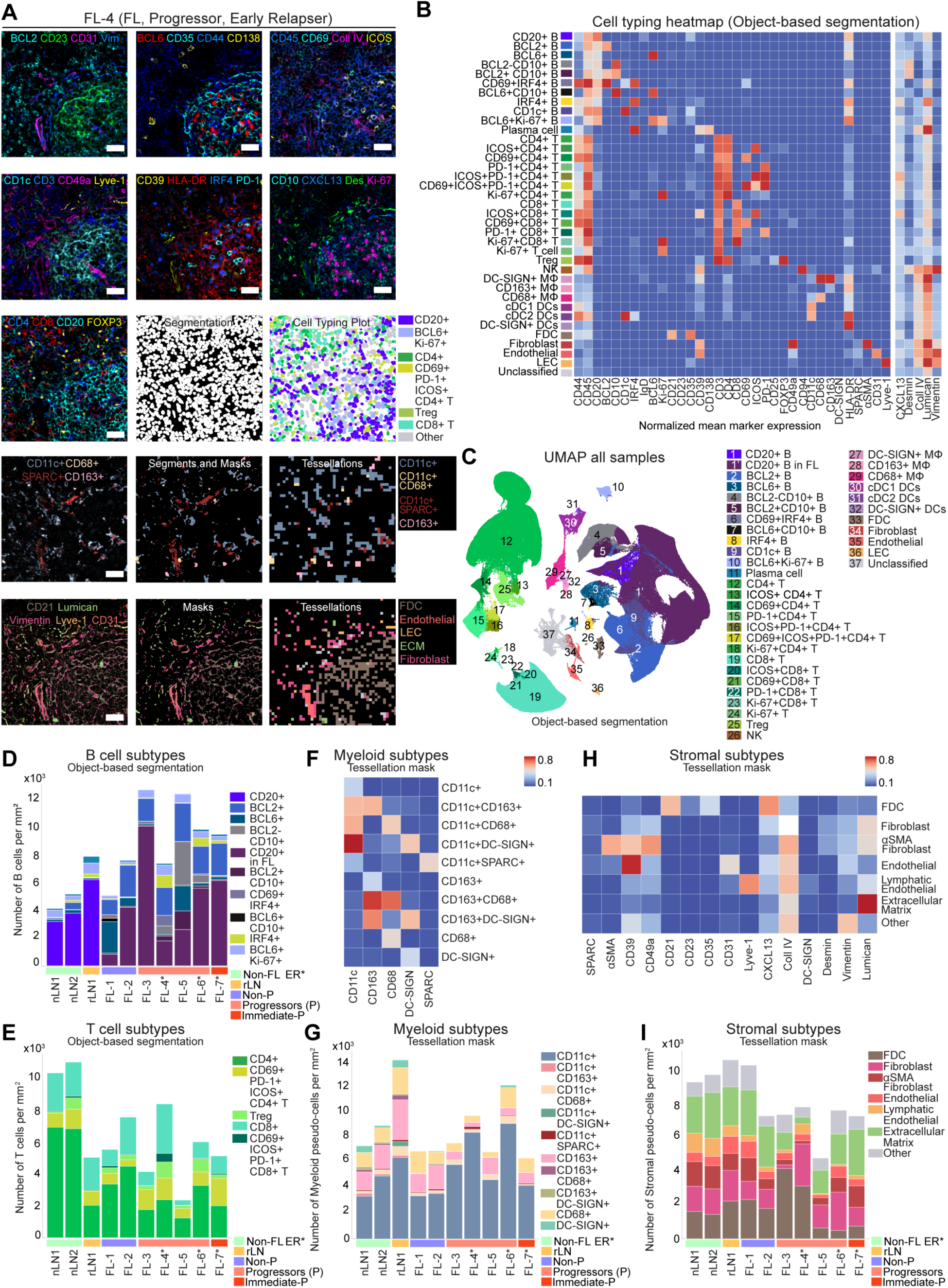
IBEX imaging provides an in-depth spatial survey of complex tissues. **(A)** Representative IBEX images of selected markers, scale bar 30 μm. Vimentin (vim), Desmin (des), Collagen IV (Coll IV). Row 3: Cell segmentation plot and cell typing plots for the same region of interest. Row 4: IBEX images of myeloid markers (left), segments and masks of immunofluorescence (IF) signal (middle), and tessellation masks of 4 myeloid populations (right). Row 5: IBEX images of stromal markers (left), segments and masks of IF images (middle), and tessellation masks of 5 stromal populations (right). **(B)** Heatmap of normalized mean marker expression used to define cell populations. 37 discrete marker clusters were identified using cell segmentation. **(C)** UMAP plot of 0.9×10^6^ cells from all samples, colored by cell populations identified by IBEX. **(D)** Quantification of major B cell subpopulations obtained from IBEX imaging and normalized by area imaged per sample. **(E)** Quantification of major T cell subpopulations obtained from IBEX imaging and normalized by area imaged per sample. **(F)** Heatmap of the normalized mean marker expression of select protein biomarkers for myeloid cell phenotyping by tessellation masks. **(G)** Quantification of major myeloid subpopulations obtained from IBEX imaging where cells are expressed as tessellation square counts per sample. **(H)** Heatmap of the normalized mean marker expression of select protein biomarkers for stromal cell phenotyping by tessellation masks. **(I)** Quantification of major stromal subpopulations obtained from IBEX imaging where cells are expressed as tessellation square counts per sample. Non-progressors (Non-P), Early relapsers (ER*).

### Cellular composition of normal and malignant follicles

Previous studies have linked clinical progression to the distribution of cell types inside and outside of B cell follicles (Farinha et al., 2010; Kridel et al., 2017). Among the samples profiled here, B cell follicles varied considerably in number and size and were identified as CD20+CD21+ structures (Figure S5A). Visual inspection and quantitative analysis revealed differences in the presence of CD8+ T cells, tingible body macrophages (CD11c+SPARC+, CD11c+CD68+) and FDCs (combinations of CD21+CD23+CD35+) in the neoplastic follicles of FL LNs (Figure 4A-B). In agreement with the literature (Börner et al., 2021; Sayin et al., 2018), the secondary follicles of non-FL LNs were enriched for GC (BCL6+Ki-67+) B cells, T follicular helper (Tfh: CD3+CD4+CD69+PD-1+ICOS+), and tingible body macrophages (Figure 4B). The early relapsers had several distinguishing patterns (Figure S5B) including the expansion of desmin+ fibroblasts around and within B cell follicles (Figure 4C). While far less abundant than other myeloid and stromal cells (Figure 4B), increased proportions of DC-SIGN+ cell subtypes were found in the follicles of early relapsers (Figure S5B), providing a potential pro-survival signal for malignant B cells via engagement with glycosylated BCRs. In contrast, DC-SIGN is traditionally found on CD163+ macrophages in the medulla and subcapsular sinus but absent from the secondary follicles of normal LNs (Figure S5B, Figure 4C). IRF4+ tumor B cells, previously implicated in aggressive FL cases with poor overall survival (Hsi et al., 2008), were identified in direct contact with DC-SIGN+ cells, Tfh cells, and cells expressing vimentin, a recognized autoantigen in FL (Cha et al., 2013) (Figure S5C). These findings are concordant with previous data demonstrating that IRF4 is upregulated in B cells following BCR engagement and/or T cell co-stimulation (Shukla and Lu, 2014). Lastly, the histological patterns identified in all three early relapsers—desmin+ FRC expansion around B cell follicles and DC-SIGN+ cells within follicles— were beginning to emerge in a non-progressor (FL-2) before therapy (Figure 4C), suggesting the possibility of incipient progression in this individual. In further support of this hypothesis, our scRNA-seq analyses revealed higher expression of the Huet gene module in this patient (FL-2) than in other early progressors (Figure 2I, Figure S1B). In summary, IBEX imaging revealed changes to the myeloid and stromal components of the TME with diagnostic and predictive potential.

**Figure 4.**
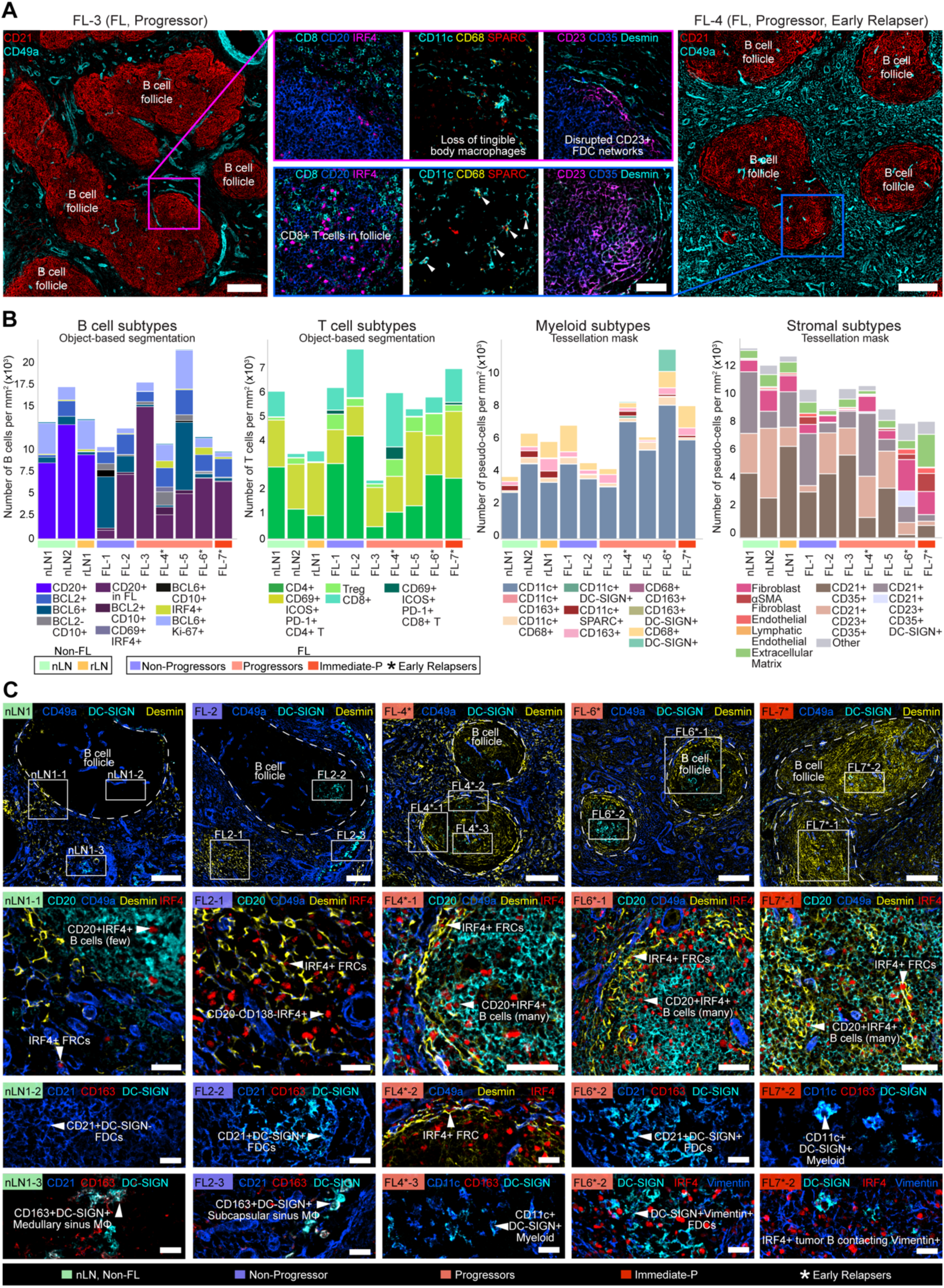
Assessing the cellular composition and histological patterns of secondary and neoplastic follicles using IBEX imaging. **(A)** IBEX images depicting differences in the shape and cellular composition of B cell follicles from FL patients, scale bars 200 μm or 50 μm (blue and magenta insets). White arrowheads denote tingible body macrophages. **(B)** Quantification of major B, T, myeloid, and stromal cells found within B cell follicles, obtained from IBEX imaging and normalized by area imaged per sample. **(C)** IBEX images depicting histological patterns shared among early relapsers: Desmin+ fibroblastic reticular cells (FRCs) inside and around B cell follicles, IRF4+ tumor B cells in contact with IRF4+ FRCs, and DC-SIGN+ myeloid or stromal cells in B cell follicles. Scale bar is 100 μm, 50 μm (Inset 1), and 25 μm (Insets 2 and 3).

### Quantitative analysis of cellular communities in diverse tissue microenvironments

Following creation of spatial maps across entire tissue sections and well-defined anatomical structures, we next evaluated the cellular ecosystems present in IBEX images. Cellular neighborhood analysis was conducted using a graph neural network-encoding approach and K-Means clustering to identify individual neighborhoods encompassing similar cell types and distributions (Goltsev et al., 2018). Cell-cell interactions were visualized on a schematic interaction graph that included 8 cell types identified by object-based segmentation, 13 stromal masks, and 5 myeloid masks using the indicated markers (Figure 5A-B). The selected cell types (8) covered major lineages identified by IBEX and scRNA-seq analysis, e.g., cycling T and Ki-67+T. The resulting approach yielded 15 neighborhoods named for the dominant cell type within each community: 5 B cell-enriched, 6 T cell-enriched, 2 myeloid-enriched, and 2 stromal-enriched (Figure 5A-B, Table S2). Importantly, this workflow classified the well-defined anatomical structures of non-FL LNs into discrete communities, e.g., GC (B1) and mantle zone (B2) (Figure 5C, Table S2). In contrast, FL LNs were highly disorganized, frequently lacking the hallmark structures of normal LNs (Figure 5D-E, Figure S5D-E, https://doi.org/10.5281/zenodo.6536724). Two B cell-enriched clusters (B1-B2) were predominantly located inside the follicles of non-FL LNs (Figure 5F, Figure S5D-E). All other B cell-enriched clusters were distributed both within and outside of the follicles (B3-B5) with B3 and B4 largely absent from non-FL LNs and expanded in certain progressors (Figure 5E, Figure S5D-E). The majority of T cell-enriched communities were located in the intrafollicular cortex and/or paracortex (T2-T6); however, one community (T1) was found inside the follicles of FL and non-FL LNs alike (Figure S5D-E). The remaining communities were myeloid (M1-M2) and stromal (S1-S2) enriched clusters corresponding to anatomical structures such as medullary and paracortical sinuses (Figure 5E-F, Table S2). We additionally evaluated the cellular neighborhoods of individual follicles using a quantitative and qualitative assessment (Figure 5G-H). The community composition of secondary and neoplastic follicles was heterogeneous across non-FL and FL samples, respectively (Figure 5G). However, individual follicles were more similar within samples than between samples, i.e., two follicles in FL-4 versus representative follicles compared between FL-4 and FL-3 (Figure 5G-H). Thus, advanced image analysis enabled identification of cellular ecosystems present in non-neoplastic LNs that were altered in malignancy.

**Figure 5.**
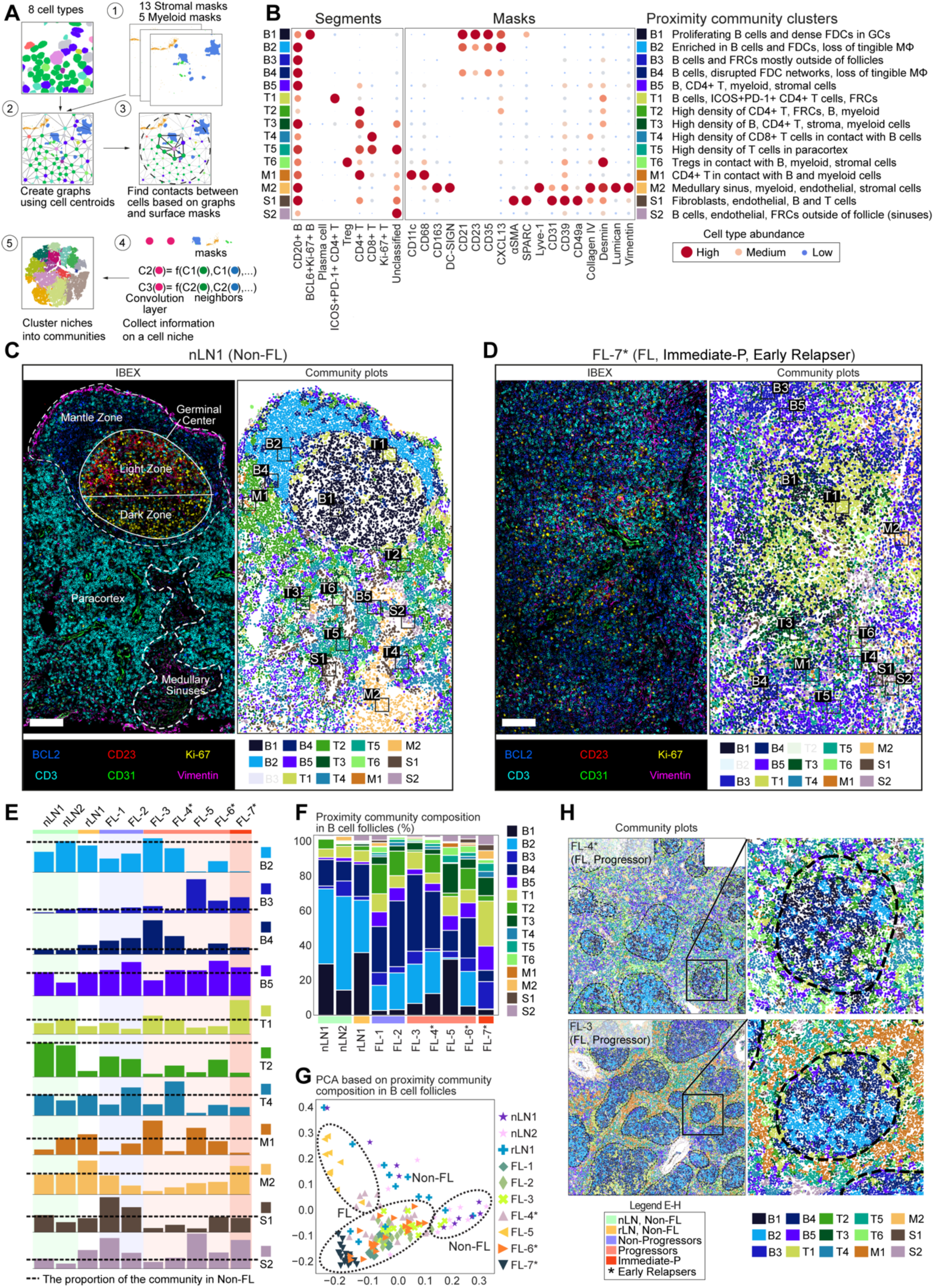
Cellular communities are shared across normal LN samples but exhibit considerable heterogeneity in tumor samples. **(A)** Overview of proximity community cluster analysis to identify cell-cell interactions using both cell segments and masks (myeloid and stromal cells). **(B)** Left: heatmap showing the relative content of cell types identified in each proximity community cluster. Right: heatmap showing the proportion of myeloid and stromal masks in proximity radius of specified cell community. Each community (one single row) contains both segmented cells from the left heat map and masks for cell types from the right heat map. **(C-D)** IBEX images depicting well organized secondary follicles in non-FL LN **(C)** and disorganized follicle in FL LN **(D)** with corresponding community plots pseudo-colored as indicated in **B**, scale bar 100 μm. **(E)** Bar plots showing the 11 most abundant proximity communities identified by IBEX for the whole imaged section. **(F)** Bar plots showing proportion of proximity communities identified by IBEX in the follicles. **(G)** Differential composition and distribution of B cell follicle communities across all normal and FL samples based on principal component analysis (PCA). **(H)** Spatial maps showing community composition of different FL samples pseudo-colored according to legend in **(B)**. Insets: community plot images of B cell follicles for selected patients.

### Quantifying unique spatial patterns across scales and imaging modalities

A dominant theme emerging from our IBEX studies was the considerable heterogeneity observed across FL samples. Because imaging of complex tissues is known to suffer from under-sampling, we extended our studies to larger tissue sections using key markers of interest. Multiplexed imaging panels, designed based on IBEX data, were applied to serial sections from FFPE samples. Following image acquisition, the conservation of histopathological patterns between IBEX small ROIs (40-plex, mean imaged area ∼9.6 mm^2^) and MxIF large ROIs (14-plex in total, 4 serial sections, mean imaged area ∼51.6 mm^2^) was evaluated (Table S2, Table S6). Samples identified to have abundant populations of follicular CD8+ T cells and DC-SIGN+ FDCs by IBEX imaging were confirmed to have these unique cell types by MxIF (Figure 6A). The presence of desmin+ stromal cells around the follicles of early relapsers and a non-progressor was additionally confirmed in FFPE tissue sections with different antibody clones (Figure 6B). Encouraged by these results, we next investigated whether follicle shape and size could determine whether the ROI selected for IBEX imaging faithfully captured the larger FL sample. While there is currently no information relating follicle heterogeneity to clinical outcome, the distinct architectural patterns we observed among FL samples (large versus small follicles, irregular versus round, abutting against one another or separate) justified this approach (Figure S6A). For quantitative assessment between samples, follicle masks were obtained through manual annotation by pathologists using CD20 and CD21 signals (Figure S5A). Using an agglomerative clustering approach, follicles were subdivided into seven subtypes and their distribution was compared between IBEX and MxIF images (Figure 6C-D). In general, IBEX and MxIF images derived from the same donor had follicles of similar shape and size (Figure 6E).

**Figure 6.**
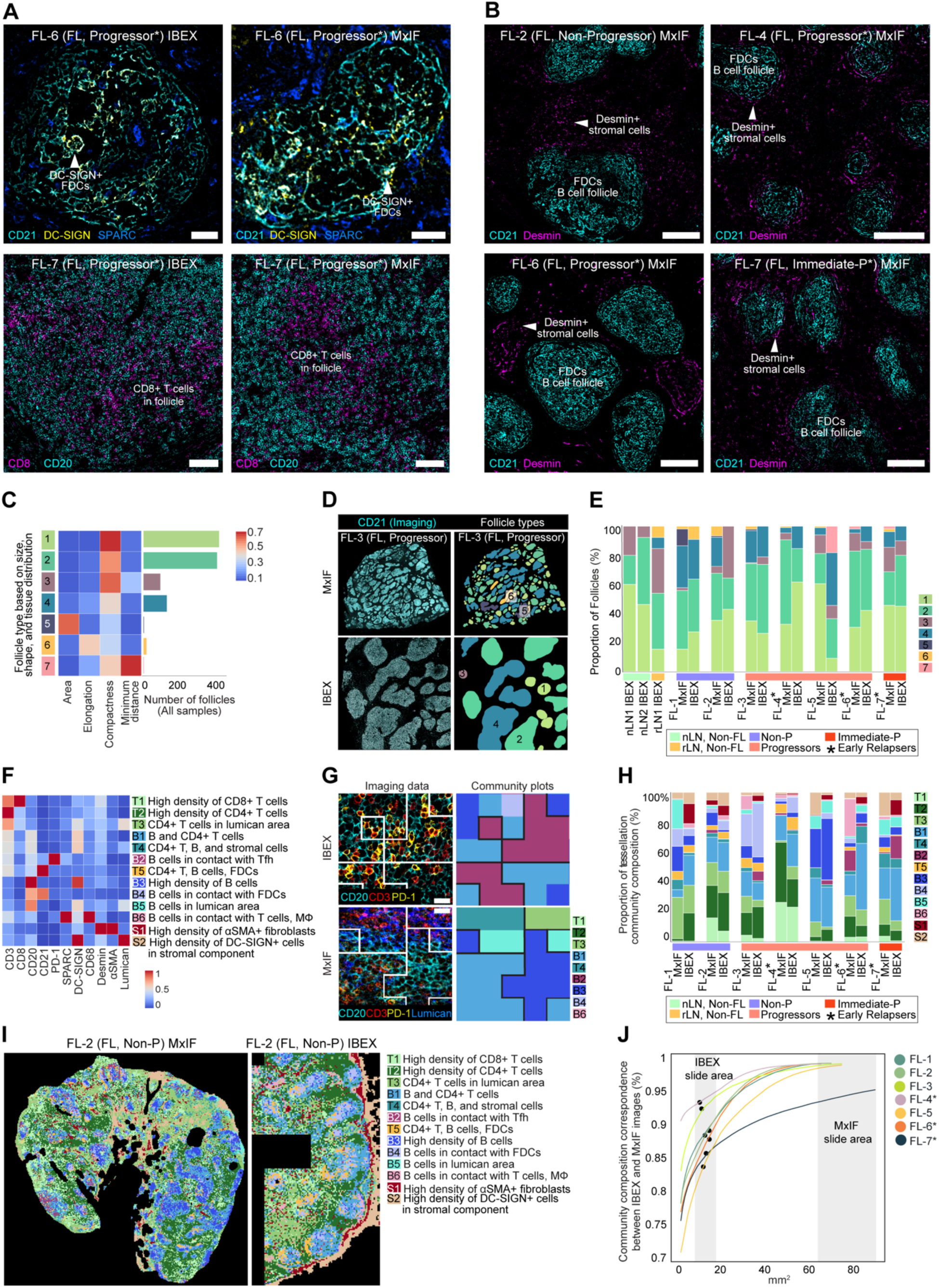
Comparison of spatial patterns and cellular communities between IBEX and MxIF images. **(A)** Confocal microscopy images from fixed frozen (IBEX) or FFPE (MxIF) samples from the same patients. Scale bar (Left, CD21 panels 50 μm; Right, CD8 panels 100 μm). **(B)** Confocal microscopy images from FFPE (MxIF) samples. Scale bar 200 μm. **(C)** Heatmap showing 7 follicle types based on shape, size, and tissue distribution (minimum distance to other follicles). Right graph: number of follicles corresponding to each cluster for all samples. **(D)** Follicle shape, size, and distribution in IBEX and MxIF images, CD21 (Cyan). Bottom row: Follicles color-coded based on types described in **(C)**. **(E)** Follicle composition of IBEX and MxIF imaged samples based on typing strategy detailed in **C**. **(F)** Heatmap of mean mask percentages per tessellation square of 11 markers measured both by IBEX and MxIF to detect tessellation community clusters. **(G)** IBEX and MxIF images with corresponding tessellation masks showing community clusters (25 μm). White lines on images (right) indicate borders of community clusters shown in adjacent plots (left). **(H)** Tessellation community plots showing the extent of correspondence between IBEX and MxIF images, color-coded based on heatmap in **F**. **(I)** Tessellation community maps for MxIF and IBEX images from one representative FL sample. **(J)** Percent similarity of IBEX and MxIF community composition depending on the area of tissue imaged and analyzed. Dots indicate the size of the IBEX region of interest.

Having observed concordance between the follicular composition of small IBEX ROIs and larger FFPE ROIs (Figure 6E), we next evaluated the community composition of these tissues using a similar approach to the one outlined in Figure 5. Following adjustment for a lower magnification objective (20X MxIF; 40X IBEX) and alignment of serial images with artifact correction (Figure S6B), tessellation masks were calculated from IBEX and MxIF images using 100 and 50 pixel-sized squares, respectively (Figure S6C). Cellular communities were defined based on 11 common markers corresponding to major cell lineages and key anatomical structures. As a result, 13 community clusters were obtained using both IBEX and MxIF data: 6 B cell-enriched clusters, 5 T cell-enriched clusters, and 2 stromal clusters (Figure 6F-G). Importantly, this community level analysis provided a qualitative and quantitative means for assessing similarities across samples and imaging modalities (Figure 6H-I). Both IBEX and MxIF images of two early relapsers (FL-6, FL-7) exhibited reduced T cell-enriched communities (T1, T2) and increased proportions of B cells in contact with CD4+ T cells (B1) (Figure 6H). To provide a metric for estimating the area of tissue to image for accurate sampling, we evaluated the mean tessellation correlation based on different sized ROIs (Figure 6J, Figure S6D). The obtained correlations were greater than 0.8, suggesting that IBEX ROIs are fairly representative of whole tissue sections.

### Integration of omics and imaging technologies for atlas creation efforts

To reach our goal of creating a reference atlas, we followed several paths to capitalize on the strengths of each technology while overcoming platform-specific limitations. First, we compared the relative abundance of major cell populations identified by bulk RNA-seq through cell deconvolution, scRNA-seq cell typing, and IBEX image analysis using cell segmentation and masks (Figure 7A). In general, similar proportions of major lymphocyte populations were observed across technologies (Figure 7A). However, myeloid and stromal cell populations were significantly underrepresented in RNA-seq datasets generated without specialized tissue dissociation methods or cell enrichment (Figure 7A). On average, IBEX images had 36 times more cells than paired scRNA-seq datasets (Table S2), empowering the study of rare cells that may require analysis of tens of thousands of cells via scRNA-seq (Figure 7B).

**Figure 7.**
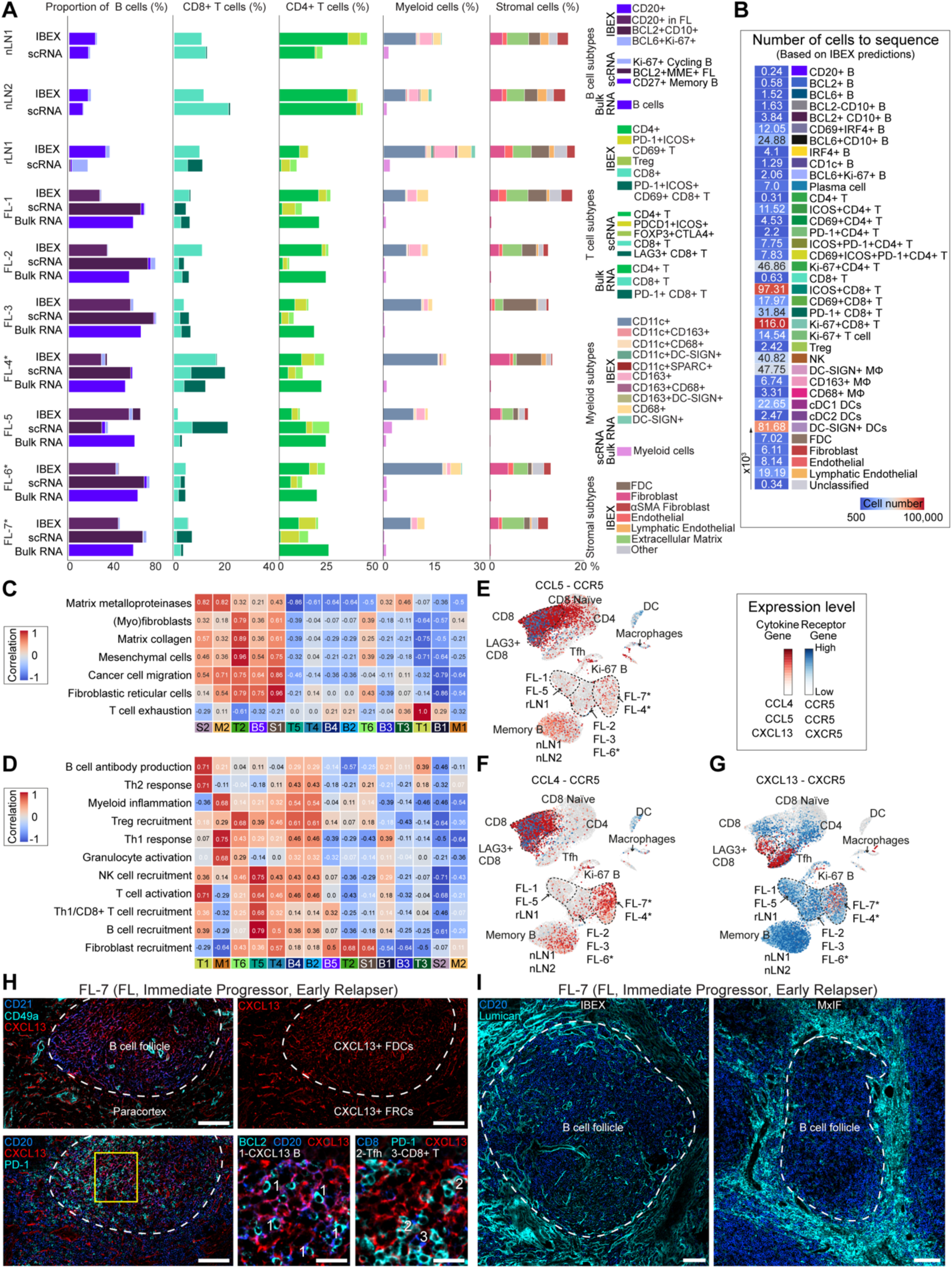
Integration of omics and imaging technologies reveals extent of stromal under-sampling and remodeling in FL TME. **(A)** Percentage of major cell populations B cells, CD4+ and CD8+ T cells, myeloid, and stromal cells measured by bulk RNA-seq (Bulk RNA), scRNA-seq (scRNA), and IBEX. Percentages for B cells, CD4+ T cells, and CD8+ T cells are compared between IBEX segmentation, scRNA-seq cell typing, and Kassandra reconstruction. Myeloid cells are compared between Kassandra reconstruction from bulk RNA-seq data and IBEX tessellation squares normalized by area of tissue imaged. Stromal cells are derived from IBEX tessellation squares normalized by area. **(B)** Heatmap showing the estimated number of cells to be profiled by scRNA-seq to identify a cluster (cell phenotype) originally identified by IBEX. Values were calculated assuming 50 cells need to be sequenced for accurate clustering and based on the average frequency of a given cell across all samples imaged by IBEX. **(C)** Heatmap showing correlations between expression of fibroblast gene signatures measured by bulk RNA-seq (Table S7) with proximity community clusters described in Figure 5. **(D)** Heatmap showing correlations between expression of cytokine gene signatures measured by bulk RNA-seq (Table S7) and proximity community clusters described in Figure 5. **(E)** UMAP of 36,212 single cells from all samples pseudo-colored for CCR5 (blue) and CCL5 (red) gene expression in the indicated cell types. **(F)** UMAP of 36,212 single cells from all samples pseudo-colored for CCR5 (blue) and CCL4 (red) gene expression. **(G)** UMAP of 36,212 single cells from all samples pseudo-colored for CXCR5 (blue) and CXCL13 (red) gene expression. **(H)** IBEX images demonstrating CXCL13+ FDCs and FRCs. Scale bar is 100 µm (large) and 20 µm (small). **(I)** IBEX and MxIF images of ECM expansion in one patient sample.

As cytokines and chemokines are essential for normal tissue organization and malignancy-induced remodeling (Nagarsheth et al., 2017), we next evaluated correlations between gene signatures curated from bulk RNA-seq with cellular communities derived from IBEX images (Figure 7C-D, Table S7, Figure S7A-B). Given gene and protein level evidence for ECM remodeling among early relapsers (Figure 2H, Figure 4C), we first explored gene signatures associated with fibrosis, including matrix metalloproteinases, collagen deposition, and (myo)fibroblasts (Figure 7C, Figure S7A). The T2 community, rich in FRCs expressing CD49a (*ITGA5*) and desmin (*Des)* (Figure S7A), was primarily found in the paracortex of non-FL LNs and the follicles of certain FL patient samples, including two early relapsers (Figure 5). Furthermore, the T1 community was correlated with gene signatures associated with B cell antibody production, Th2 immunity, and T cell exhaustion (Figure 7C-D, Table S7). These molecular features are in line with the cellular composition (PD-1+ Tfh cells, B cells, and CXCL13+ FRCs) and expansion of the T1 community in the follicles of an early relapser (FL-7) (Figure 5).

In addition to these fibroblast-associated gene signatures, bulk RNA-seq identified cytokines strongly correlated with IBEX communities, including the major pro-fibrotic factor TGF-β and its isoforms (Figure S7A-B, Table S7) (Luzina et al., 2015; Prud’homme, 2007). Several chemokines (CCL4, CCL5, CXCL13) involved in the recruitment of diverse cell types to the TME were also found (Table S7). Our scRNA-seq analysis revealed elevated CCL4 and the CXCL13-CXCR5 gene pair in FL B cells from two early relapsers (FL-4, FL-7) (Figure 7E-G). LAG3+ CD8+ T cells, present only in rLN1 and FL LNs (Figure 2F), showed elevated levels of CCL4-CCR5, CCL5-CCR5, and CXCL13-CXCR5 gene pairs (Figure 7E-G). Increased numbers of CD8+ T cells infiltrated the follicles of two early relapsers (FL-4, FL-7, Figure 4A-B, Figure 6A), suggesting a potential mechanism involving tumor B cell recruitment of CD8+ T cells through the secretion of CCL4 and other soluble factors. We used IBEX imaging to confirm the presence of CXCL13+ FL B cells, described before (Husson et al., 2002), and expansion of CXCL13+ FDCs and CD49a+ FRCs in FL patients, also described before (Mourcin et al., 2021) (Figure 7H). Lastly, we evaluated the extent of stromal desmoplasia (collagen deposition, ECM remodeling) in these samples. Dense networks of lumican+ fibers were found around the follicles of early relapsers (Figure 7I, Figure S7C), likely providing a pauci-cellular or acellular boundary around malignant B cells.

## DISCUSSION

Atlases constructed from even a modest number of samples, such as outlined here, represent critical resources for discovery efforts and translational research (Börner et al., 2021; Conde et al., 2022; Eraslan et al., 2022; Jones et al., 2022; Liu and Zhang, 2022; Regev et al., 2017; Rozenblatt-Rosen et al., 2020; Snyder et al., 2019). Accordingly, our multi-scale, multimodal approach yielded several insights and technical advances. One advance is our detailed annotation of the diverse cell types and anatomical structures present in normal and malignant LNs. Another key finding is the identification of several distinguishing features in the TMEs of early relapsers, the most high-risk of FL patients. These included the upregulation of *FOXP1* by tumor B cells, a prognostic biomarker for adverse response to immunochemotherapy in FL patients (Mottok et al., 2018). The malignant B cells of early relapsers exhibited several characteristics consistent with antigen engagement within the TME, e.g., high levels of IRF4 protein and upregulation of BCR signaling pathways. We also report the expansion of desmin+ fibroblasts and ECM deposition around the follicles of patients who relapsed within 2 years of frontline therapy. The histological patterns we identified in 40-plex images were confirmed in clinically relevant FFPE samples using whole slide imaging with a reduced marker set. In addition to highlighting IBEX as a potential biomarker discovery platform, this work suggests that antibodies directed against FOXP1, IRF4, desmin, vimentin, lumican, and DC-SIGN may warrant inclusion in established diagnostic panels for risk-adapted management of FL patients. To this end, immunohistochemical (IHC) evaluation of intratumoral vimentin was recently identified to predict histologic transformation in FL patients (Madsen et al., 2019) and IHC of FOXP1 has been proposed as a low cost screen for high risk patients (Mottok et al., 2018).

Using NGS and advanced imaging technologies, we profiled cell suspensions via bulk RNA-seq, 36,212 cells via scRNA-seq, and 1.8×10^6^ cells via object-based segmentation of IBEX images from non-FL donors and untreated FL patients prospectively enrolled in a clinical trial. Three patients were later identified as early relapsers; however, the biopsies described here were collected prior to any therapeutic intervention. Through the application of advanced deconvolution (bulk RNA-seq) and clustering (RNA-seq and IBEX) algorithms, cell populations were annotated into distinct cell types: 10 using bulk sequencing data, 22 employing scRNA-seq methods, and 37 based on IBEX with object-based segmentation. Myeloid and stromal cells pose several challenges for accurate quantification including the requirement for enzymatic tissue disassociation and inaccurate cellular segmentation for single cell and spatial assays, respectively (Germain et al., 2022; Gerner et al., 2012; Risom et al., 2022). For this reason, we developed a workflow that phenotyped diverse myeloid and stromal populations using pixel-based tessellation masks in intact tissues. Imaging studies, 95.9 mm^2^ (IBEX, 10 samples) and 361.1 mm^2^ (MxIF, 7 samples) in total area, provided additional insight into the location and cell-cell interactions of diverse cell types. To empower future atlas studies, we have made these datasets publicly available for exploration using the ASCT+B Reporter (https://hubmapconsortium.github.io/ccf-asct-reporter/) (Börner et al., 2021) (https://doi.org/10.5281/zenodo.6536724).

Bulk RNA-seq has provided insight into the mutational burden, copy number alterations, and cellular composition of various tumors (Rozenblatt-Rosen et al., 2020). Additional studies characterized genomic aberrations and transcriptomic differences in FL B cells shown to correlate with progression (Dave et al., 2004; Huet et al., 2018; Pastore et al., 2015). The cost-effectiveness of bulk RNA-seq, in combination with previously reported mutations and disease-associated gene expression signatures, make this an attractive approach for stratifying patients for personalized treatment options. Here, we extend the utility of bulk analysis by correlating gene signatures associated with cell types and states to cellular communities found *in situ*. Despite these advances, our results demonstrate that bulk RNA-seq data does not provide the same level of detailed information on the gene expression profiles necessary for molecular dissection of rare subpopulations of cells (Hwang et al., 2018; Rozenblatt-Rosen et al., 2020).

To resolve heterogenous expression patterns at single cell resolution, we additionally profiled normal and FL LNs using scRNA-seq. Importantly, the data we collected from normal mesenteric LNs may contribute to a contemporary map of the human lymphatic system, one of the last frontiers of anatomical medicine. Our scRNA-seq studies also revealed the existence of multiple subclones in FL patients with dominant clones emerging in clinical cases associated with early progression and relapse after therapy. Others have speculated on the critical role for antigen selection in the clonal evolution of FL B cells (Bahler et al., 1992). Given previous studies demonstrating spontaneous apoptosis of isolated FL cells *in vitro,* along with the identification of self-reactive tumor cells in FL patients (Cha et al., 2013; Sachen et al., 2012), a compelling argument can be made for TME-derived autoantigens driving positive selection of malignant clones. Within the TME, T cells have been shown to express multiple immune checkpoint inhibitors in scRNA-seq datasets derived from FL patients (Andor et al., 2019). We also observed the enrichment of CD8+ T cells expressing several markers associated with T cell dysfunction and/or exhaustion including *PDCD1 (PD-1), LAG3*, *HAVCR2* (*TIM3*), *and CTLA4* (Armengol et al., 2021). Interestingly, these CD8+ T cells also expressed molecules related to cytotoxic T cell function such as *PRF1*, *GZMA*, and *NKG7* (Ng et al., 2020; Szabo et al., 2019). The co-expression of markers associated with cytolytic activity and immune inhibition suggests the presence of an anti-FL response that might be re-invigorated for therapeutic purposes. Although initial efforts using T-cell directed cell therapies in FL have shown great promise, including among high-risk patients, mechanisms of immune evasion and resistance remain largely unknown (Fowler et al., 2022).

We also evaluated the composition of normal and FL LNs using a high content imaging method. IBEX spatially resolved myeloid, stromal, and rare (<0.05%) cells that were completely absent or significantly undercounted in RNA-seq datasets. Using 19 myeloid and stromal-enriched antibody targets, in combination with analytical methods optimized for typing irregular shaped cells, we quantified diverse myeloid and stromal subsets within major and minor anatomical structures of the LN. In single tissue sections, we visualized changes to the stromal and myeloid components of the FL TME including loss of FDC meshworks, reduction in tingible body macrophages, cell-cell contacts between DC-SIGN+ cells and FL B cells, and expansion of CD49a+ lymphoid stromal cells (Carbone et al., 2019; Kridel et al., 2017; Mourcin et al., 2021; Verdière et al., 2018). The ability to phenotype diverse cell populations across spatial scales—whole tissue sections, major anatomical structures, proximity communities—presents an opportunity to resolve the confusing literature concerning cellular distribution patterns and clinical outcome in FL (Glas et al., 2007; Kridel et al., 2017; Verdière et al., 2018). Granular phenotyping with high content imaging will aid these efforts by 1) distinguishing between tumor promoting and tumor suppressing cell types, 2) identifying not one but a series of spatial patterns contributing to progression, and 3) interrogating the cellular ecosystems of FL B cells.

Although IBEX is fully compatible with FFPE samples (Radtke et al., 2022), the construction of multiplexed imaging panels is a time and resource intensive process (Hickey et al., 2021). An additional consideration is the total time required to image large >10 mm^2^ regions of interest with cyclic imaging methods such as IBEX. To overcome these challenges, we performed MxIF on serial FFPE sections using 4 distinct panels comprised of antibodies directed against key biomarkers identified via IBEX. Notably, this approach provided a means to validate unique spatial patterns, such as DC-SIGN+ FDCs, using an orthogonal method and alternative antibody clones. As an additional resource, we devised analytical approaches to compare morphological features shared between IBEX and MxIF images prepared from the same patient. Here, the shape, size, and distance between B cell follicles yielded a unique structural fingerprint for assessing the similarities between small and large ROIs. Following alignment of serial sections into one composite image, we evaluated communities shared between IBEX and MxIF images. The resulting workflow offers a means to 1) identify key spatial patterns and biomarkers of interest using high content imaging, 2) apply predictive markers to diagnostic samples, 3) compare samples acquired through complementary platforms, and 4) calculate the area required to adequately profile a sample. This last point is especially relevant given the growing need for operator-independent strategies for accurate sampling of heterogeneous TMEs (Berry et al., 2021).

A challenge for tumor atlas efforts is to go beyond a detailed description of tissues to a greater understanding of how genetic alterations and spatial patterns contribute to pathogenesis, clonal evolution, and treatment response. To achieve this aim, multimodal data must be fully integrated. Here, we highlight several biological features enriched in FL patients who progressed early and relapsed shortly after therapy. Using scRNA-seq, B cells from early relapsers were distinguished for their significant upregulation of pathways involved in BCR signaling, cytokine signaling, and immune activation. IBEX imaging revealed high numbers of IRF4+ FL B cells in direct contact with DC-SIGN+ and vimentin+ cells in the TMEs of these patients, suggestive of active signaling and BCR engagement through endogenous lectins and/or autoantigens, respectively (Amin et al., 2015; Carbone et al., 2019; Cha et al., 2013; Shukla and Lu, 2014). FL B cells are known to engage in a bidirectional crosstalk with lymphoid stromal cells, including cytokine/chemokine-producing fibroblasts (Mourcin et al., 2021). We hypothesize that these interactions, coupled with matrisome-associated factors produced by FL B cells, results in the expansion and fibrogenic potential of desmin+ fibroblasts found around the follicles of early relapsers. Bulk RNA-seq and IBEX community analysis confirmed the expression of fibroblast and cytokine genes, e.g., TGF-β, known to impact ECM deposition and functional remodeling of LN tissues. As with other cancers (Risom et al., 2022), stromal desmoplasia was shown to be a distinctive feature in this patient cohort, distinguishing early relapsers from other FL patients based on further evaluation of FFPE tissue sections with MxIF imaging. Advanced fibrotic lesions are often poorly vascularized and characterized for the dense physical barriers they form, two properties that can significantly impair drug delivery (Pakshir et al., 2020). Together, these findings strongly encourage careful examination of anti-fibrotic agents, coupled with therapies that blunt BCR signaling, for the treatment of FL patients who relapse after frontline therapy.

In summary, we present a comprehensive molecular and spatial atlas of normal and malignant LNs taken from untreated FL patients in the context of a prospective clinical trial. In so doing, we have addressed several key questions in the field including: What unique information is provided by bulk RNA-seq, scRNA-seq, and multiplexed tissue imaging? How can we effectively integrate these datasets to leverage the strengths of each technology? How extensively are myeloid and stromal cells under-sampled by sequencing approaches? Most importantly, this work presents the unique opportunity to profile the TMEs of FL patients prior to therapeutic intervention. Despite the urgent clinical need, there are several impediments to the careful examination of relapsed FL including its broad clinical and genetic heterogeneity and challenges with recruiting sufficient numbers of patients with adequate tissue biopsies for large studies (Rodgers and Barr, 2021). Therefore, this work may inform the selection of novel therapeutic approaches for early relapsers, the highest priority in follicular lymphoma clinical trials (Maddocks et al., 2017). The extension of these studies to a greater number of samples with divergent clinical outcomes will only highlight the value of this approach by identifying predictive biomarkers and histological patterns of early disease progression and treatment resistance.

## LIMITATIONS OF THE STUDY

The primary limitation of this study is the modest number of samples evaluated using single biopsies collected from different sites of the human body. Non-neoplastic LNs are known to exhibit anatomical and functional differences depending on their site of drainage (Grant et al., 2020). While challenging to implement, the intra-tumor heterogeneity observed among FL patients provides significant rationale for multi-site profiling (Araf et al., 2018; Haebe et al., 2021). An additional constraint is the lack of paired bulk RNA-seq and MxIF datasets from non-FL LNs due to technical challenges with performing paired analyses from these small LNs. As we did not employ dissociation or enrichment protocols for single cell analysis of myeloid and stromal cells, these populations are significantly underrepresented in our single cell datasets as compared to other studies that explicitly examined these populations (Buechler et al., 2021; Mourcin et al., 2021). Finally, extensive domain expertise is needed to carefully annotate the vast diversity of cell types present in these samples, especially cells of the stromal lineage that are known to vary across tissues and disease states (Pradhan et al., 2021). We anticipate these classifications will evolve with technologies and additional multiomic studies arising from atlas efforts.

## ACKNOWLEDGMENTS

This research was supported by the Intramural Research Program of the NIH, NIAID and NCI. Z.Y. and B.C.L. are supported by the BCBB Support Services Contract HHSN316201300006W/HHSN27200002 to Medical Science & Computing, LLC. D.J. is supported by the grant of the European Research Council (ERC); European Consolidator Grant, XHale (Reference #771883). M.C.K is supported by Frederick National Laboratory for Cancer Research contract 75N91019D00024. We are deeply appreciative of Arlene Radtke, Anita Gola, Ellen Quardokus, Rebecca Beuschel, and Derek Einhaus for their advice, careful review of materials, and support.

## AUTHOR CONTRIBUTIONS STATEMENT

A.J.R. wrote the manuscript. A.J.R., E.P., and A.B. directed the analysis and interpretation of the data. A.L.S., L.M.S., W.H.W., M.R., and R.N.G. conceived the study and contributed to the writing of the manuscript. A.J.R. acquired, processed, and assisted with analysis and annotation of IBEX and MxIF imaging data. A.V., V.S., I.G., P.O., E.P., M.S., M.P., and G.P., conducted IBEX and MxIF data processing and analysis. A.J.R., E.P., E.J., and S.P performed histological assessment of the tissue slides. S.I., E.S., Y.L., A.L.S., M.C.K., and D.W.H. executed and analyzed scRNA-seq experiments. A.B., O.K., M.M., E. N., N.K. and performed bulk RNA-seq and WES data processing. Z.R.Y. and B.L. developed image alignment software and registered all IBEX images. A.J.R., E.P., A.V., E.S., A.B., L.Y., M.S., O.K., A.B., M.M., M.P., I.G., V.S., S.I., G.P., Y.L., A.L.S., P.O., K.N., E. N., N.K., and R.A. designed display items. L.Y. designed Figure 1. A.J.R. annotated scRNA-seq and IBEX data using agreed upon Cell Ontology terms. S.P. and T.D.H. prepared samples for multiomic assessment and performed diagnostic pathology on FL samples. D.J., T.D.H., E.J., S.P, J.D.P, J.M., N.F., J.L.D., and J.M.H provided technical insight, reagents, and tissues. All authors participated in the final review of the text.

## METHODS

### Clinical study details

All patients were enrolled on a prospective clonal evolution study for adults with grade I-II or 3A follicular lymphoma who have not received systemic therapy and are without evidence of histologic transformation [NCT03190928]. All patients consented to the trial. The primary endpoint of the study is time to initiation of frontline systemic therapy. The primary objective is to analyze the molecular biology of patients with progression within 2 years of study entry (‘early progressors’) compared to patients who do not progress and need therapy within 2 years of study entry (‘non-progressors’). The secondary objective is to characterize the molecular biology of patients who relapse <2 years after frontline therapy (‘early relapsers’). Baseline staging procedures include computed tomography (CT) and fluorodeoxyglucose (FDG)-positron emission tomography (PET) scans along with bone marrow biopsy with aspirate, and patients are staged by the Lugano criteria (Cheson et al., 2014). All patients are offered excisional LN biopsy, if feasible. Enrolled patients are assigned a baseline FLIPI score (Solal-Celigny et al., 2004) and initially assessed by uniform protocol-defined treatment criteria to determine need for immediate frontline therapy. For those who do not meet criteria for treatment, they are monitored with clinic visits every 4 months for 2 years, every 6 months in years 3-5, and then annually until they meet criteria for treatment. CT scans are every 8 months for 2 years, then annually. FDG-PET scans are repeated at 2 years and any time of suspected progression. Normal human mesenteric LNs were obtained from patients undergoing elective risk-reducing gastrectomies or colon resections for colon adenocarcinoma at the National Cancer Institute (NCI) based on an Institutional Review Board (IRB) approved tissue collection protocol (#13C-0076). Biopsies of these LNs were grossly normal as determined by the operative surgeon and histopathologically normal as determined by an expert pathologist. Follicular hyperplasia was identified in one sample (rLN1) based on standard diagnostic evaluation with hematoxylin and eosin (H&E). See Table S1 for clinical and pathological characteristics of patients. *BCL2* rearrangements were identified using fluorescence *in situ* hybridization (FISH), the gold standard for translocation detection in lymphoma FFPE (Allahyar et al., 2021). Representative sections from all samples were evaluated using H&E staining before multimodal analysis.

### Sample preparation from human tissues

The size of the LNs ranged from less than 1 cm (nLN1, nLN2), 2 cm (rLN1), to 6 cm (FL1-7) in diameter. Unfixed LN were measured (L x W x H), cut along the longitudinal axis, and macroscopically inspected. For routine assessment, the LN was sectioned into slices <4mm in thickness and prepared as FFPE samples as previously described (Jonigk et al., 2011). Depending on the size of the LN, samples were additionally prepared as snap frozen tissue blocks (< 5mm^3^) and cell suspensions (at least 20% of total LN volume). All clinical stains were performed with automated immunostainers, BenchMark Ultra (Roche) or BOND-Max (Leica Biosystems), according to the manufacturers’ instructions. Diagnostic panels consisted of CD20 (clone L26), CD10 (clone SP67), CD3 (clone 2GV6), BCL2 (clone SP66), BCL6 (clone EP278), Ki-67 (clone MIB-1), CD21 (clone EP3093), CD23 (clone IB12), and IgD (clone 92).

For bulk and scRNA-seq, cell suspensions were prepared by manual disruption of the tissues and frozen down viably. Following tissue homogenization, cells were frozen and stored at -150°C in liquid nitrogen. Prior to sequencing, single cell suspensions were thawed rapidly in a 37°C water bath until ice had just disappeared, then transferred to a 50 ml tube and washed with 50 ml of cold (4°C) 1xPBS. Viable cells were enumerated manually using trypan blue exclusion. For IBEX imaging, human LNs (1 cm^3^ or smaller in size) were fixed with BD CytoFix/CytoPerm (BD Biosciences) diluted in PBS (1:4) for 2 days. Following fixation, all tissues were washed briefly (5 minutes per wash) in PBS and incubated in 30% sucrose for 2 days before embedding in OCT compound (Tissue-Tek) as described previously (Radtke et al., 2022; Radtke et al., 2020). Non-FL LNs (nLN1, nLN2, rLN1) were only analyzed by scRNA-seq and IBEX imaging.

### Whole exome sequencing (WES) analysis

Low quality reads were filtered using FilterByTile/BBMap v37.90 (Bushnell, 2014) and aligned to human reference genome GRCh38 (GRCh38.d1.vd1 assembly) using BWA v0.7.17 (Li and Durbin, 2009). Duplicate reads were removed using Picard’s v2.6.0 MarkDuplicates (“Picard Toolkit”, 2019. Broad Institute, GitHub Repository. http://broadinstitute.github.io/picard/; Broad Institute). Indels were realigned by IndelRealigner and recalibrated by BaseRecalibrator and ApplyBQSR using tools taken from GATK v3.8.1 (Auwera and O’Connor, 2020). Somatic single nucleotide variations (sSNVs), small insertions, and deletions were all detected using Strelka v2.9 (Saunders et al., 2012).

### Bulk RNA-seq processing and analyses

RNA was isolated from cell suspensions using the AllPrep kit (Qiagen) and libraries were generated using the TruSeq Stranded mRNA Library kit (Illumina). Paired end sequencing was performed on an Illumina NextSeq2000. RNA-seq reads were aligned using Kallisto v0.42.4 to GENCODE v23 transcripts 69 with default parameters. The protein-coding transcripts, immunoglobulin heavy, kappa and lambda light chains, and TCR-related transcripts were retained. Noncoding RNA, histone, and mitochondria-related transcripts were removed, resulting in 20,062 protein coding genes. Gene expression was quantified as the sum of the transcripts and re-normalized per million (TPM) and log2-transformed (Goldman et al., 2020).

### Deconvolution of bulk RNA-seq

The Kassandra machine learning algorithm was used to predict cell percentages from bulk RNA-seq (Zaytcev et al., 2020). The model consisted of a two-level hierarchical ensemble that used LightGBM as building blocks. The model was trained on artificial RNA-seq mixtures of different cell types (T cells, B cells, NK, macrophages, cancer-associated fibroblasts, and endothelial cells) obtained from multiple datasets of sorted cells. All datasets were isolated from poly-A or total RNA-seq profiled human tissues with read lengths higher than 31 bp and at least 4 million coding read counts. These datasets passed quality control by FASTQC with minimal contamination (< 2%). The model was trained to predict the percentage of RNA belonging to specific cell types. Predicted percentages of RNA were later converted into percentages of cells using the methodology described previously (Racle et al., 2017). For Figure 2, Figure 7C-D, and Figure S7A-B, gene signature scores were calculated using the ssGSEA algorithm from the GSVA R package (Hänzelmann et al., 2013). Raw scores were medium scaled to (−2, 2) or to (−3, 3) range. See Tables S2-S4.

### scRNA-seq processing and analysis

Viable cells were diluted in 1xPBS such that when loaded on the 10x Genomics Chromium Controller they were at a capture number of ∼6,000 cells. After capture, single cell RNA-seq/VDJ libraries were generated using the 10X Chromium Single Cell 5’ gene expression/V(D)J kit and processed according to the manufacturer’s instructions. Sequencing of libraries was performed on an Illumina NOVA-Seq and cycling was performed according to the manufacturer’s suggestions. All samples had captures performed on the same day with reagents from the same kit. All were sequenced together on multiple sequencing runs to achieve target depth.

FAST-Q files were processed through the 10X Cell Ranger Pipeline v5.0.1 with alignment to a GRCh38 reference (refdata-gex-GRCh38-2020-A for gene expression reads and refdata-cellranger-vdj-GRCh38-alts-ensembl-5.0.0 for TCR and BCR enriched reads). A classical scRNA-seq analysis pipeline was performed as described in (Luecken and Theis, 2019). The Cell Ranger 4.0 tool and Scanpy 1.7.1 package with Python 3.6 were used (Wolf et al., 2018). Cells with less than 1,000 unique molecular identifiers (UMIs), more than 10,000 UMIs, and cells with more than 10% mitochondrial gene UMIs were removed. These criteria were empirically selected after initial data analysis in Cell Ranger. These thresholds were applied to minimize several technical clusters (small number of UMIs, high mitochondrial gene expression, and lack of meaningful marker genes). Selection of the top 3,000 over-dispersed genes was performed as described (Stuart et al., 2019). Next, log-transformation of the data and linear regression of expression data against the number of UMIs and number of genes in cells was performed followed by kNN graph construction. The Leiden algorithm was used for cell clustering with the top 3,000 marker genes and not the whole transcriptome. Elbow plot analysis was used to determine how many PCs were needed to capture the majority of the variation in the data (Traag et al., 2019). Data were visualized with UMAP. After overall analysis of all datasets, we selected the subset of T cells, B cells, and other cells and performed an analysis of each subset. Cell types were identified based on marker gene expression. Moreover, cells from the same cell type were selected based on the Leiden clusters and not the UMAP coordinates.

MIXCR v.2.1.7 (Bolotin et al., 2015) was used to analyze BCR sequences from RNA-seq data. Single clonotypes were grouped into clones with unique VDJ combinations and identical CDR3 nucleotide sequences. B cell clones were further aggregated into clone groups if the VDJ combination was the same and if the CDR3 nucleotide sequences differed no more than 1 nucleotide (Figure S2A). For definition of CDRs and *de novo N*-linked glycosylation sites (Figure S2B), BCR sequences, heavy and light chain, from scRNA-seq data were identified by Loupe VDJ Browser analysis and assigned to individual single cells. Predominant BCR clonotypes were further analyzed by porting the amino acid sequences into abYsis (http://www.abysis.org/abysis/).

Gene set enrichment analysis was performed using the C2CP gene set from the mSigDB (Liberzon et al., 2015). Analysis was performed in R v4.1.3 using fgsea v1.20.0 (Korotkevich et al., 2021). Analysis was performed on a ranked gene set resulting from the log fold change values from differential expression analysis using the FindMarkers function in Seurat v4.1.0 (Hao et al., 2021). Ribosomal genes were removed before analysis was performed. Plots were generated in ggplot2 v3.3.5 (Wickham, 2016). Dot plots were generated using the DotPlot function in Seurat. Gene module scores for individual cells were generated using the AddModule function in Seurat, with the number of control features set to the same length as the gene set of interest. Gene modules were visualized using the VlnPlot function in Seurat. For Figure 2G-H, the B cell populations included cycling B, tumor B, GC B, memory B, naïve B, and FCRL4+ B single cell clusters.

### High content imaging using IBEX

High content imaging was performed on fixed frozen sections as described previously (Radtke et al., 2022; Radtke et al., 2020). Briefly, 20 μm sections were cut on a CM1950 cryostat (Leica) and adhered to 2 well Chambered Coverglasses (Lab-tek) coated with 15 μl of chrome alum gelatin (Newcomer Supply) per well. Frozen sections were permeabilized, blocked, and stained in PBS containing 0.3% Triton X-100 (Sigma-Aldrich), 1% bovine serum albumin (Sigma-Aldrich), and 1% human Fc block (BD Biosciences). Immunolabeling was performed with the PELCO BioWave Pro 36500-230 microwave equipped with a PELCO SteadyTemp Pro 50062 Thermoelectric Recirculating Chiller (Ted Pella) using a 2-1-2-1-2-1-2-1-2 program. A complete list of antibodies and an IBEX LN antibody panel can be found in Table S6. Cell nuclei were visualized with Hoechst (Biotium) and sections were mounted using Fluoromount G (Southern Biotech). Mounting media was thoroughly removed by washing with PBS after image acquisition and before chemical bleaching of fluorophores. After each staining and imaging cycle, samples were treated for 15 minutes with 1 mg/mL of LiBH_4_ (STREM Chemicals) prepared in diH_2_O to bleach all fluorophores except Hoechst and Alexa Fluor 594.

### Whole slide MxIF imaging of FFPE tissue sections

5 µm tissue sections were cut from FFPE samples and placed onto glass slides. Prior to immunolabeling, tissue sections were baked in a 60°C oven for 1 hour to adhere the tissues to the slides. Deparaffinization was performed with 2 exchanges of 100% xylene (10 minutes per exchange) followed by 100% ethanol for 10 minutes, 95% ethanol for 10 minutes, 70% ethanol for 5 minutes, and 10% formalin for 15 minutes. Antigen retrieval was performed by incubating slides in AR6 buffer (Akoya Biosciences) for 40 minutes in a 95°C water bath. After 40 minutes, slides were removed from the water bath and allowed to cool on the bench for 20 minutes. Blocking and immunolabeling was performed using the PELCO BioWave Pro 36500-230 microwave according to the steps outlined in Table S6. Prior to immunolabeling, tissue sections were outlined with an ImmEdge pen to create a hydrophobic barrier (Vector laboratories) and then rehydrated with PBS. Following a 30-minute incubation in blocking buffer, tissue sections were incubated with primary antibodies, washed 3 times in PBS, and then incubated with appropriate secondary antibodies. Directly conjugated primary antibodies were applied last after blocking with 5% normal rabbit and/or goat sera (Abcam). Cell nuclei were visualized with Hoechst (Biotium) and sections were mounted using Fluoromount G (Southern Biotech).

### Image acquisition and alignment

Representative sections from different tissues were acquired using an inverted Leica TCS SP8 X confocal microscope equipped with 20X (NA 0.75) and 40X objectives (NA 1.3), 4 HyD and 1 PMT detectors, a white light laser that produces a continuous spectral output between 470 and 670 nm as well as 405, 685, and 730 nm lasers. Panels consisted of antibodies conjugated to the following fluorophores and dyes: Hoechst, Alexa Fluor (AF)488, AF532, phycoerythrin (PE), PE/iFluor594, eF570, AF555, AF594, iFluor (iF)594, AF647, eF660, AF680, and AF700. All images were captured at an 8-bit depth, with a line average of 3, and 1024×1024 format with the following pixel dimensions: x (0.284 μm, 40X), y (0.284 μm, 40X), x (0.568 μm, 20X), y (0.568 μm, 20X), and z (1 μm). Images were tiled and merged using the LAS X Navigator software (LAS X 3.5.5.19976). For IBEX tissue imaging, multiple tissue sections were examined before selecting a representative tissue section that contained several distinct follicles, often resulting in unusually shaped region of interests. For multiplexed imaging of FFPE tissue sections, whole tissue sections were imaged using a 20X objective. To ensure proper alignment over distinct imaging cycles, careful attention was paid to the quality of image stitching achieved with the Leica software and z-stacks were set by manual inspection of notable features such as unusually shaped nuclei throughout the tissue volume. These unusual features were matched across the z-stack and over multiple cycles of IBEX as outlined in a detailed protocol (Radtke et al., 2022). Fluorophore emission was collected on separate detectors with sequential laser excitation of compatible fluorophores (3-4 per sequential) used to minimize spectral spillover. The Channel Dye Separation module within the LAS X 3.5.5.19976 (Leica) was then used to correct for any residual spillover. For publication quality images, gaussian filters, brightness/contrast adjustments, and channel masks were applied uniformly to all images. Image alignment of all IBEX panels was performed as described previously (Radtke et al., 2022; Radtke et al., 2020) using SimpleITK (Lowekamp et al., 2013; Yaniv et al., 2018). Software can be downloaded via a zip file from the Imaris extensions code repository (https://github.com/niaid/imaris_extensions/archive/refs/heads/main.zip). Installation instructions are available online: [https://github.com/niaid/imaris_extensions] and in the README.md file which is part of the zip file. Additional details can be found in the XTRegisterSameChannel SimpleITK Imaris Python Extension YouTube tutorial (https://youtu.be/rrCajI8jroE). Please see sample data on Zenodo for usage of the software [https://doi.org/10.5281/zenodo.4632320].

To obtain multiplexed images of FFPE tissue sections, serial sections were imaged with 4 distinct panels of antibodies containing Hoechst and 2-8 antibodies per panel (Table S6). Following image acquisition, images were aligned using Hoechst as a fiducial. As a first approximation, a pathologist manually obtained rotation matrices for cell-cell alignment across serial sections using downsampled Hoechst channels and GIMP (GNU Image Manipulation Program) image editor. Following image rotation for alignment, the register_translation function from the skimage python package was used to find a translation vector. Upon identification of a suitable rotation matrix and translation vector based on Hoechst+ nuclei, these parameters were applied to all serial image stacks. The register_translation function uses cross-correlation in Fourier space, optionally employing an upsampled matrix-multiplication DFT (Discrete Fourier transform) to achieve arbitrary subpixel precision (Guizar-Sicairos et al., 2008). Artifacts (such as fluorophore aggregates and uneven staining) were manually masked by 3 pathologists.

### IBEX image analysis: Object-based segmentation

Object-based cellular segmentation was performed using a convolutional neural network (CNN)-based approach with Mask R-CNN architecture and ResNet-50 as a backbone (He et al., 2017). As an input three channels were used: Hoechst for nuclei, CD45 as a base membrane, and composite of several other membrane markers (CD138, CD163, CD94, CD69, CD8, CD4). Composite images were made by choosing the brightest pixel across all marker channels for each separate pixel position. Final images were normalized to a range of 0-1 by dividing by the maximal possible intensity value (255). The prediction window size was set to 256 pixels on each side. To stitch prediction tiles, the original images were cropped into intersecting windows with 136 pixel steps. A 60 pixel buffer was used on every side of prediction window (256 - 60 * 2) for predicted cell selection in overlapping areas. Buffer size was chosen as a half of a maximum potentially possible cell side size. The training settings for the deep learning algorithms used in this work are given in Table S5.

### IBEX image analysis: Cell typing

Among the 39 antibodies included in the multiplexed imaging panels, three distinct staining patterns were observed: membrane or cytoplasmic (all markers except for BCL6, Ki-67, FOXP3, IRF4, CD68), nuclear (BCL6, Ki-67, FOXP3, IRF4), and endosomal/lysosomal (CD68). The CNNs were trained to recognize the presence or absence of a biomarker based on these patterns of expression for individual cells. The input image consisted of three channels: Hoechst, marker of interest, and segment mask for a cell of interest. Input images were cropped to 128×128 pixels, corresponding to the cell bounding box and its nearest environment, and normalized to values ranging from 0-1, dividing by the maximal possible intensity value. All three networks had the same architecture (ResNet-50) with two neurons in the last layer (signal present/signal absent) with Softmax activation function in the output layer. The Softmax activation function allows normalization of the CNN’s output to values from 0 to 1, where 1 corresponds to positive expression, 0 corresponds to negative expression of a given marker, and 0.5 corresponds to an uncertain prediction (hard case for NN, outcomes have equal weights). This approach is also sensitive to the spatial distribution of each specific marker. Every possible ground truth cell type can be expressed as a sequence of markers that must be present (encoded as 1), may be present (creating two versions of this cell type, with 0 and 1) and must be absent (encoded as 0). For example, a particular cell may be defined as a B cell if this cell is CD20+ (encoded as 1), CD21+/- (encoded as 1 or 0), and CD3- (encoded as 0)). Cosine similarity is calculated between every sequence of cell expression and every cell type encoding vector. The final cell type is defined as the closest match based on probability. Cell annotations and counts across samples can be found in Table S2.

### Tessellation masks for myeloid and stromal cells

Masks for certain immune, myeloid, and stromal markers were generated by Otsu’s method (Otsu, 1979). The following markers were masked for IBEX images: CD11c, CD21, CD23, CD31, CD35, CD39, CD49a, CD68, CD163, CXCL13, Collagen IV, DC-SIGN, Desmin, Lumican, Lyve-1, SPARC, α-SMA, and Vimentin. The following markers were masked for MxIF images: CD3, CD8, CD20, CD21, CD68, DC-SIGN, Desmin, Lumican, PD-1, SPARC, and α-SMA. All channel masks were inspected by pathologists and compared to the raw immunofluorescence signal. In a few select instances of poor algorithm performance, thresholds were manually corrected using Fiji. We define a tessellation as a process of splitting a mask derived from a marker of interest into non-intersecting squares that covers a full image area. A signal density heatmap is then created by computing the percentage of positive pixels in each of these squares. Every square can be roughly considered as a “pseudo-cell” measuring 16×16 pixels for stromal and myeloid subpopulations identified in IBEX images. Percentages of masks in each square “pseudo-cell” is equivalent to mean cell marker expression for object-based cellular segmentation.

For phenotyping stromal cells from IBEX imaging data, tessellation-based analysis was performed with over a dozen markers (CD21, CD23, CD31, CD35, CD39, CD49a, CXCL13, Collagen IV, Desmin, Lumican, Lyve-1, SPARC, α-SMA, and Vimentin), including several markers not exclusive to stromal cells (e.g. CD39, CD49a, SPARC). For markers expressed by stromal and non-stromal elements, e.g. CD39 on Tregs, stromal masks were first created to mark an area of interest based on co-localization with lineage-defining markers, e.g. CD31 for endothelial cells and vimentin for mesenchymal cells. For well-described stromal markers (CD31, Lyve-1, CD21, Desmin, Vimentin, Lumican), tessellation masks were created directly from the channel data and thresholded by 0.2, an empirically derived parameter. To elaborate, this limit (0.2) corresponded to the value where the amount of ‘masked pixels’ and ‘empty pixels’ were equal and intersected on an x-y plot with the ‘mask threshold’ on the x-axis and ‘fraction of pixels covered by mask’ on the y-axis (Figure S3D). Clustering was then performed on masked pixel data using the unsupervised clustering algorithm Phenograph (Levine et al., 2015) with cosine distance metric and 30 nearest neighbors. For phenotyping myeloid cells from IBEX imaging data, a similar approach was implemented using 5 markers (CD11c, CD163, CD68, DC-SIGN, and SPARC) where only the SPARC marker was not exclusive to macrophages and dendritic cells. Although CD1c and HLA-DR were detected on myeloid cell subtypes, these markers are also expressed on B cell subpopulations. Due to contaminating signal from B cells, the dominant population in FL samples, CD1c and HLA-DR were excluded from myeloid cell phenotyping. For CD68 and CD163 markers, the tessellated mask was an integration of pixel-level data and corresponding macrophage segment masks. To exclude empty tessellation squares, we filtered empty squares by a 0.1 threshold. A different masking strategy was applied for myeloid cells in order to detect the pseudopodia of these cell types which typically cover less area than stroma. K-Means was used as a clustering algorithm. These tessellation-based approaches were also used to define cellular communities present in both IBEX and MxIF images using 100 and 50 pixel-sized squares, respectively. For these analyses we used 11 markers and the K-Means clustering algorithm. Cell annotations and counts across samples can be found in Table S2.

### Follicles shape analysis of IBEX and MxIF images

To compare the follicle shape across small and large ROIs acquired using distinct imaging methods and sample preparations (IBEX/fixed frozen versus MxIF/FFPE), masks were applied to B cell follicles and sample parameters were calculated. B cell follicle masks from IBEX images were obtained by manual annotation from pathologists based on concentrated areas positive for CD20 and CD21 and reduced lumican signal for most samples (Figure S5A). Follicle masks were created based on CD21 and CD20 signal alone for MxIF images. Once masks were generated, the area, minimal distance to the nearest follicle, elongation, and compactness were calculated for each follicle. As a final step agglomerative clustering with ward linkage was performed on given follicle shape parameters (Ward Jr, 1963). The following elongation formula was used:

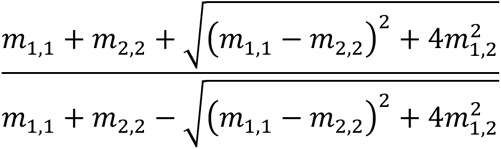

Compactness formula where A and P stand for area and perimeter correspondingly: 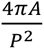.

### Community analysis of IBEX images

To perform community analysis, a cell neighborhood graph was generated with Delaunay triangulation algorithm from SciPy spatial algorithms package using coordinates derived from cell centroids (Virtanen et al., 2020). In this graph, each cell is represented as a node, and adjacent cells are connected via edges. To remove outlier edges, we aggregated lengths of all edges for all cells in the dataset across all samples, and a threshold value was selected as the 95th percentile of all lengths – 28.4 µm, edges longer than this threshold were removed. After this operation, we assigned features for every node in the graph, which used this model: 1) cell type assigned as categorically encoded vector, 2) percentages of selected binary masks in 56.8 µm radii underneath this cell, 3) median distance of edges connected to this node. Using this graph, an Adversarially Regularized Variational Graph Auto-Encoder (Fey and Lenssen, 2019; Pan et al., 2018) was trained to obtain short descriptive vector representation of cell neighborhoods in an unsupervised, generative-adversarial manner. The framework consists of two models - a variational graph autoencoder and an adversarial model (3 layered perceptron). The latter model is used for regularization of the main graph autoencoder model. During training, the autoencoder learns to correctly predict node edges and reject randomly added non present edges. In parallel the discriminator model is trained to distinguish between learned autoencoder representations and their random permutations, thus regularizing the model to closely follow the latent data distribution, since the autoencoder is rewarded for correct generation of data, close to the real distribution. Thus, in the process of training, an autoencoder model learns representation of tissue topology, since it can reconstruct correct contacts for a given cell and has learned representations that closely follow the true data distribution. These representations, or embeddings, learned by the autoencoder, allow clustering and detection of cell similarities by topological features of the given tissue.

The model was trained for a fixed number of epochs (100) and the model with lowest loss was selected for downstream analysis. The trained autoencoder predicted embedding vectors for all samples in the cohort and these vectors were then clustered using the K-means algorithm to obtain 15 different clusters, which represent neighborhoods of cells or communities. To facilitate community description, the mean cell composition and mean mask percentages were calculated for each community. Communities were grouped based on the dominant cell type and morphological structures present in them: B- and T- cells enriched neighborhoods, myeloid- and stroma- enriched neighborhoods. Communities were visualized by drawing cell contours and coloring them according to community type. A detailed description of IBEX communities (Figure 5) and comparison with MxIF communities (Figure 6) can be found in Table S2. The training settings for the deep learning algorithms used in this work are given in Table S5.

### Slide concordance analysis

Tessellation-based communities were used to analyze the concordance of small IBEX ROIs with large, full tissue section ROIs from MxIF. For each sample, we performed sampling with window side sizes varying from 2,500 to 17,000 pixels (2.02-98.8 mm^2^) with 500 pixel steps (284 µm). Crops were sampled uniformly, with distances between centers equaling 200, 300, 400, and 500 pixels for sides of 2500-4000, 4500-6000, 6500-8500, and 10000-17000 pixels correspondingly. Crops with a tissue area of less than 50% were excluded from further analysis. We then measured the Pearson correlation between the percentage of tessellation communities for given crops and the full slide. Graphics displays mean correlation for given crop size.

### Predicting the number of cells to be sequenced based on IBEX data

For Figure 7B, we predict the number of cells that need to be profiled by scRNA-seq based on frequencies obtained from IBEX imaging data. Cell numbers were calculated based on cell frequencies obtained from the entire 1.8×10^6^ cell IBEX dataset. A cluster size of 50 cells was used to estimate the number of cells to be sequenced.

### Correlation analysis between RNA-seq and IBEX communities

For Figure 7C-D and Figure S7A-B, fibroblast and cytokine gene signatures were manually curated from the literature (see Table S7). Pairwise correlations were performed between RNA-seq gene signatures, described in *Deconvolution of bulk RNA-seq,* and IBEX communities, described in *Community analysis of IBEX images*.

## LIFE SCIENCE TABLE WITH EXAMPLES FOR AUTHOR REFERENCE

| REAGENT or RESOURCE | SOURCE | IDENTIFIER |
| --- | --- | --- |
| Antibodies |  |  |
| CD20 (clone L26) | Ventana Medical Systems | Cat#760-2531;<br>RRID:<br>AB_2335956 |
| CD10 (clone SP67) | Ventana Medical Systems | Cat#790-4506;<br>RRID:<br>AB_2336021 |
| CD3 (clone 2GV6) | Ventana Medical Systems | Cat#790-4341;<br>RRID:<br>AB_2335978 |
| BCL2 (clone SP66) | Ventana Medical Systems | Cat#790-4604;<br>RRID: NA |
| BCL6 (clone EP278) | Cell Marque | Cat#227R-28;<br>RRID: NA |
| Ki-67 (clone MIB-1) | Agilent | Cat#M7240; RRID:<br>AB_2142367 |
| CD21 (clone EP3093) | Ventana Medical Systems | Cat#760-4438;<br>RRID: NA |
| CD23 (clone IB12) | Leica Biosystems | Cat# NCL-CD23-1B12; RRID:<br>AB_442058 |
| IgD (clone 92) | Agilent | Cat# A0093; RRID:<br>NA |
| CD20 AF488 (clone L26) | Thermo | Cat#53-0202-82;<br>RRID:<br>AB_10734358 |
| CD20 eF660 (clone L26) | Thermo | Cat#50-0202-82;<br>RRID:<br>AB_11150959 |
| SPARC AF532 (goat polyclonal, custom conjugate from company) | R&D | Cat#AF941; RRID:<br>AB_2892754 |
| CD10 PE (clone FR4D11) | Caprico Biotechnologies | Cat#103926;<br>RRID: NA |
| CD10 PE (clone HI10a) | BioLegend | Cat#312204;<br>RRID: AB_314915 |
| CD3 AF594 (clone UCHT1) | BioLegend | Cat#300446;<br>RRID:<br>AB_2563236 |
| BCL2 AF647(clone 100) | BioLegend | Cat#658705;<br>RRID:<br>AB_2563279 |
| Collagen IV (rabbit polyclonal) | Abcam | Cat#Ab6586;<br>RRID: AB_305584 |
| Goat anti-rabbit IgG AF700 | Thermo | Cat#A21038;<br>RRID:<br>AB_2535709 |
| IgD AF488 (clone IA6-2) | BioLegend | Cat#348216;<br>RRID:<br>AB_11150595 |
| CD21 AF532 (clone Bu32), custom conjugate from company | BioLegend | Cat#NA; RRID:<br>AB_2892739 |
| CD138 PE (clone MI15) | BioLegend | Cat#356504;<br>RRID:<br>AB_2561878 |
| BCL6 AF647 (clone K112-91) | BD Biosciences | Cat#561525;<br>RRID:<br>AB_10898007 |
| CD31 AF700 (clone WM59) | BioLegend | Cat#303133;<br>RRID:<br>AB_2566326 |
| HLA-DR AF488 (clone L243) | BioLegend | Cat#307620;<br>RRID: AB_493175 |
| CD23 AF532 (clone EBVCS-5), custom conjugate from company | BioLegend | Cat#NA; RRID:<br>AB_2892740 |
| CD1c PE (clone L161) | BioLegend | Cat#331506;<br>RRID:<br>AB_1088999 |
| CD163 AF647 (clone GH1/61) | BioLegend | Cat#333620;<br>RRID:<br>AB_2563475 |
| CD11c AF700 (clone B-Ly6) | BD Biosciences | Cat#561352;<br>RRID:<br>AB_10612006 |
| CD8 AF488 (clone SK1) | BioLegend | Cat#344716;<br>RRID:<br>AB_10549301 |
| CD4 AF532 (clone RPA-T4) | Thermo | Cat#58-0049-42;<br>RRID:<br>AB_2802361 |
| FOXP3 eF570 (clone 236A/E7) | Thermo | Cat#41-4777-82;<br>RRID:<br>AB_2573609 |
| CD25 AF647 (clone M-A251) | BioLegend | Cat#356128;<br>RRID:<br>AB_2563588 |
| Ki-67 AF700 (clone B56) | BD Biosciences | Cat#561277;<br>RRID:<br>AB_10611571 |
| ICOS AF488 (clone CS98.4A) | BioLegend | Cat#313514;<br>RRID:<br>AB_2122584 |
| SPARC AF532 (goat polyclonal, custom conjugate from company) | R&D | Cat#AF801; RRID:<br>AB_2892755 |
| PD-1 PE (clone EH12.2H7) | BioLegend | Cat#329906;<br>RRID: AB_940483 |
| CD69 AF647 (clone FN50) | BioLegend | Cat#310918;<br>RRID: AB_528871 |
| CD39 FITC (clone A1) | BioLegend | Cat#328206;<br>RRID: AB_940425 |
| Lyve-1 AF532 (goat polyclonal, custom conjugate from company) | R&D | Cat#AF2089;<br>RRID:<br>AB_2892756 |
| CD35 PE (clone E11) | BioLegend | Cat#333406;<br>RRID:<br>AB_2292231 |
| CD68 AF647 (clone KP1) | Santa Cruz | Cat#sc-20060;<br>RRID:<br>AB_2891106 |
| a-SMA AF488 (clone 1A4) | Thermo | Cat#53-9760-82;<br>RRID:<br>AB_2574461 |
| a-SMA eF660 (clone 1A4) | Thermo | Cat#50-9760-82;<br>RRID:<br>AB_2574362 |
| Lumican AF532 (goat polyclonal, custom conjugate from company) | R&D | Cat#AF2846;<br>RRID:<br>AB_2892757 |
| IRF4 PE (clone IRF4.3E4) | BioLegend | Cat#646404;<br>RRID:<br>AB_2563005 |
| DC-SIGN AF647 (clone 9E9A8) | BioLegend | Cat#330112;<br>RRID:<br>AB_1186092 |
| Desmin AF488 (clone Y66) | Abcam | Cat# Ab185033;<br>RRID:<br>AB_2892748 |
| CD49a AF647 (clone TS2/7) | BioLegend | Cat#328304;<br>RRID:<br>AB_1236407 |
| CD94 AF488 (clone DX22) | BioLegend | Cat#305506;<br>RRID: AB_314536 |
| Vimentin AF532 (clone O91D3) custom conjugate from company | BioLegend | Cat#NA; RRID:<br>AB_2892753 |
| CD45 PE/iFluor594 (clone F10-89-4) | Caprico Biotechnologies | Cat#1016185;<br>RRID: 2892742 |
| CD44 AF647 (clone IM7) | BioLegend | Cat#103018;<br>RRID: AB_493681 |
| BCL2 (clone SP66) | Abcam | Cat# Ab236221;<br>RRID: NA |
| Donkey anti-rabbit IgG AF594 | Thermo | Cat#A-21207;<br>RRID: AB_141637 |
| CD10 (polyclonal) | R&D | Cat#AF1182;<br>RRID: AB_354652 |
| Donkey anti-goat IgG AF680 | Thermo | Cat#A-21084;<br>RRID: AB_141494 |
| CD21 (clone SP186) | Abcam | Cat#Ab240987;<br>RRID: NA |
| Donkey anti-rabbit IgG AF555 | Thermo | Cat#A-31572;<br>RRID: AB_162543 |
| CD68 iFluor594 (clone KP1) | Caprico Biotechnologies | Cat#1064135;<br>RRID: 2892745 |
| DC-SIGN (clone h209) | LS Bio | Cat#LS-B3782;<br>RRID:<br>AB_10689801 |
| Donkey anti-rat IgG AF647 | Jackson ImmunoResearch | Cat#712-605-153;<br>RRID:<br>AB_2340694 |
| SPARC (polyclonal) | R&D | Cat#AF941; RRID:<br>AB_355728 |
| HI-6B Multiplex Panel - Human CD3, CD4, CD8, FoxP3 | Cell IDx | Cat#HI06B-005 |
| CD3 (clone SP7) | Abcam | Cat#Ab16669;<br>RRID: AB_443425 |
| Goat anti-rabbit IgG AF532 | Thermo | Cat#A-11009;<br>RRID:<br>AB_2534076 |
| PD-1 (polyclonal) | Novus | Cat#AF1086;<br>RRID: AB_354588 |
| Donkey anti-goat IgG AF555 | Thermo | Cat#A-21432;<br>RRID:<br>AB_2535853 |
| Chemicals, peptides, and recombinant proteins |  |  |
| Trypan Blue Exclusion | Thermo | 15250061 |
| Triton X-100 | Sigma-Aldrich | Cat#T8787 |
| Tween 20 | Millipore Sigma | Cat#9005-64-5 |
| PBS, pH 7.4 | GIBCO | Cat#10010-023 |
| BD Cytotfix/Cytoperm | BD Biosciences | Cat#554722 |
| Optimal cutting temperature (OCT) compound | Sakura | Cat#4583 |
| Sucrose | Millipore Sigma | Cat#S0389 |
| Bovine Serum Albumin | Millipore Sigma | Cat#A1933 |
| Human Fc-block | BD Biosciences | Cat#564219 |
| diH<sub>2</sub>O | Quality Biological | Cat#351-029-101 |
| Fluoromount-G | Southern Biotech | Cat#0100-01 |
| Hoechst 33342 | ThermoFisher Scientific | Cat#H3570 |
| Lithium borohydride (purchase in 1 gram aliquots) | STREM Chemicals | Cat#93-0397 |
| Chrome Alum Gelatin | Newcomer Supply | Cat#1033A |
| AR6 buffer 10X | Akoya Biosciences | Cat#AR600250ML |
| Ethanol, 200 Proof | Decon Labs, Inc. | Cat#2701 |
| Formalin, 10% neutral buffered | Cancer Diagnostics, Inc. | Cat#FX1003 |
| Xylene, histology grade | Newcomer Supply | Cat#1446C |
| ImmEdge Pen | Vector Laboratories | Cat#H-4000 |
| Normal Rabbit Serum | Abcam | Cat#Ab7487 |
| Normal Goat Serum | Abcam | Cat#Ab138478 |
| Critical commercial assays |  |  |
| Chromium Next GEM Single Cell 5' Kit v2, 16 rxns | 10X Genomics | Cat#PN-1000263 |
| Chromium Next GEM Chip K Single Cell Kit, 48 rxns | 10X Genomics | Cat#PN-1000286 |
| Chromium Single Cell Human BCR Amplification Kit, 16 rxns | 10X Genomics | Cat#PN-1000253 |
| Library Construction Kit, 16 rxns | 10X Genomics | Cat#PN-1000190 |
| AllPrep kit | Qiagen | Cat# 80204 |
| TruSeq Stranded mRNA Library kit | Illumina | Cat# 20020594 |
| Deposited data |  |  |
| RNA-seq data | This paper | TBD |
| Imaging data (IBEX and MxIF) | This paper | TBD |
| ASCT+B Tables | This paper | TBD |
| Software and algorithms |  |  |
| Leica Application Suite X (LAS X) | Leica Microsystems | RRID:SCR_013673 |
| Imaris and Imaris File Converter (x64, version 9.5.0) | Bitplane | RRID:SCR_007370 |
| Python (version 3.7.0 and higher) | Python | RRID:SCR_008394 |
| SimpleITK Imaris Python Extension | (Radtke et al., 2022) | https://doi.org/10.5281/zenodo.4632320 |
| BostonGene Software | This paper | TBD |
| GraphPad Prism, version 8 | GraphPad Software | RRID:SCR_002798 |
| Adobe Photoshop CC 2020 | Adobe |  |
| Adobe Illustrator CC 2020 | Adobe |  |
| Adobe After Effects CC 2020 | Adobe |  |
| Adobe Media Encoder CC 2020 | Adobe |  |
| Other |  |  |
| 2-well chambered coverglass | Lab-Tek | Cat#155380 |
| Dissecting mat, flexible, polypropylene | Newcomer Supply | Cat#5218A |
| Dissecting needles | Newcomer Supply | Cat#5220PL |
| Histomolds, 15mm x 15mm x 5mm | Sakura | Cat#4566 |
| Sterile disposable scalpels #11 | Newcomer Supply | Cat#6802A |
| VWR Superfrost Plus micro slides | VWR | Cat#48311-703 |
| EasyDip slide staining kit | Newcomer Supply | Cat#5300KIT |
| EasyDip anodized aluminum jar rack holder | Newcomer Supply | Cat#5300JRK |
| Wash N'Dry cover slip rack | Electron Microscopy Sciences | Cat#70366-16 |

## SUPPLEMENTAL INFORMATION TITLES AND LEGENDS

### Contents

Figure S1: Evaluation of gene expression profile and cellular composition of normal and FL LNs using scRNA-seq.

Figure S2: RNA-seq assessment of clonotypes and *N*-glycosylation sites within variable regions of immunoglobulin genes.

Figure S3. Artificial intelligence (AI)-driven pipeline for multiplexed image analysis.

Figure S4: IBEX imaging allows detailed spatial profiling of tumor B, myeloid, stromal, and other cell types *in situ*.

Figure S5. Visualization and quantification of histological patterns and cellular communities using IBEX.

Figure S6. Workflows for comparing spatial patterns between IBEX and MxIF images.

Figure S7. Integration of multimodal technologies to evaluate normal and FL LNs.

Table S1. Clinical and pathological characteristics of non-FL and FL samples.

Table S2. Comparison of samples across omics and imaging platforms (Included as separate dataset).

Table S3. Differentially expressed genes from bulk RNA-seq reported as fold change values (Included as separate dataset).

Table S4. Cell-specific gene expression and cell annotations for bulk RNA-seq based deconvolution.

Table S5. Training settings for deep learning algorithms used in this study.

Table S6. IBEX and MxIF imaging panels for fixed frozen and FFPE tissues.

Table S7. Bulk RNA-seq gene signatures used for cell community assessment. Accompanying Datasets: https://doi.org/10.5281/zenodo.6536724

