## Supplemental Information for "A Multi-scale, Multiomic Atlas of Human Normal and Follicular Lymphoma Lymph Nodes"

**Contents:**

Figure S1: Evaluation of gene expression profile and cellular composition of normal and FL LNs using scRNA-seq.
Figure S2: RNA-seq assessment of clonotypes and *N*-glycosylation sites within variable regions of immunoglobulin genes.
Figure S3. Artificial intelligence (AI)-driven pipeline for multiplexed image analysis.
Figure S4: IBEX imaging allows detailed spatial profiling of tumor B, myeloid, stromal, and other cell types *in situ*.
Figure S5. Visualization and quantification of histological patterns and cellular communities using IBEX.
Figure S6. Workflows for comparing spatial patterns between IBEX and MxIF images.
Figure S7. Integration of multimodal technologies to evaluate normal and FL LNs.

Table S1. Clinical and pathological characteristics of non-FL and FL samples.
Table S2. Comparison of samples across omics and imaging platforms (Included as separate dataset).
Table S3. Differentially expressed genes from bulk RNA-seq reported as fold change values (Included as separate dataset).
Table S4. Cell-specific gene expression and cell annotations for bulk RNA-seq based deconvolution.
Table S5. Training settings for deep learning algorithms used in this study.
Table S6. IBEX and MxIF imaging panels for fixed frozen and FFPE tissues.
Table S7. Bulk RNA-seq gene signatures used for cell community assessment.

Accompanying Datasets: https://doi.org/10.5281/zenodo.6536724

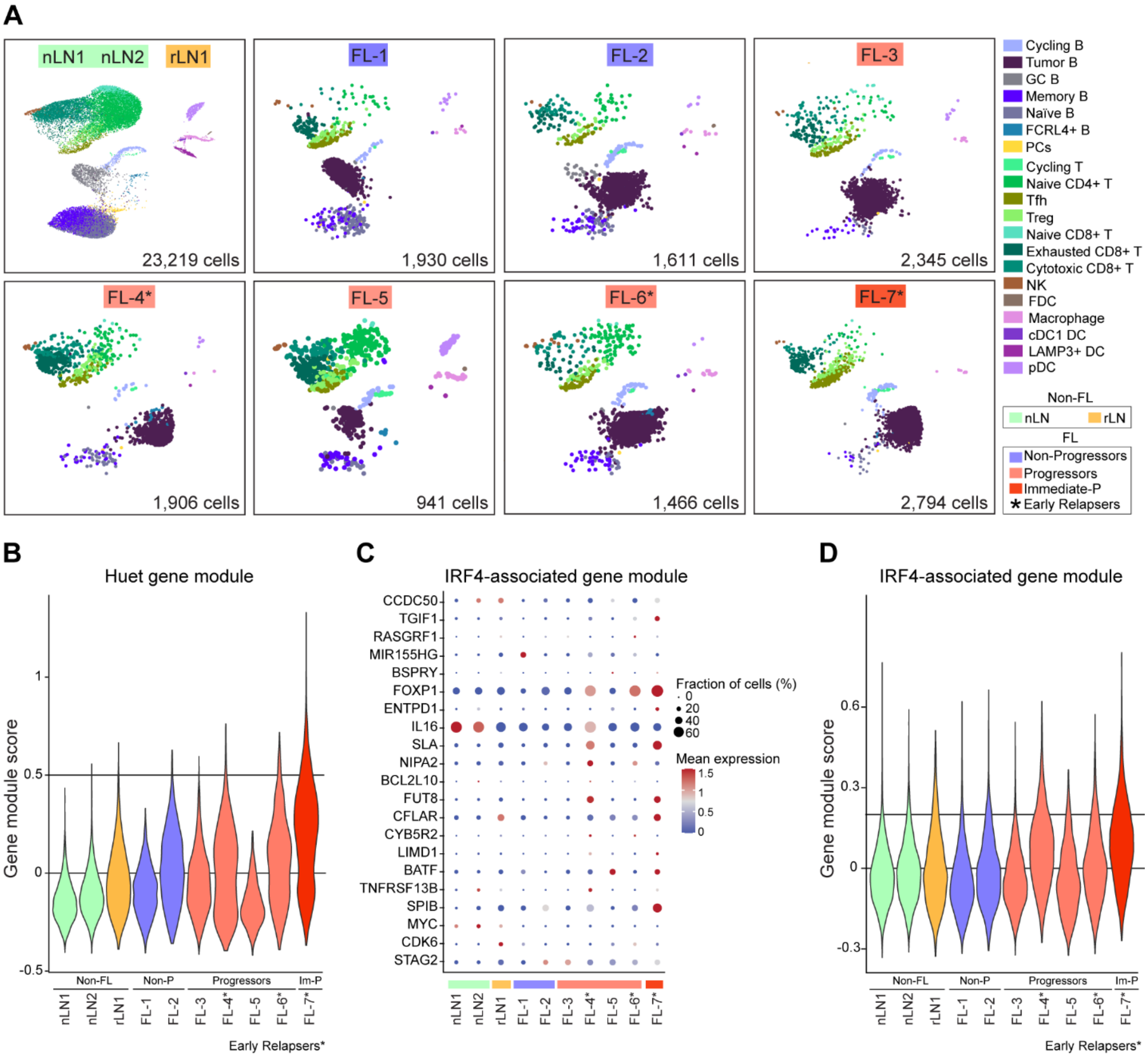


**Figure S1: Evaluation of gene expression profiles and cellular composition of normal and FL LNs using scRNA-seq.**

**(A)** UMAP plots from individual samples colored by cell populations measured by scRNA-seq. Cells from non-FL samples are included in one plot. **(B)** Violin plot of Huet gene module for all samples. *PDRM15* and *SMISA8* were not included due to no expression from the 23 Huet gene signature (Huet et al., 2018). See Figure 2I for individual genes. **(C)** Dot plot depicting dynamic expression of individual genes associated with IRF4 expression in hematological malignancies (Wang et al., 2014). **(D)** Violin plot of IRF4-associated gene module for all samples. For B and D, horizontal lines are included to aid in visual comparison only.

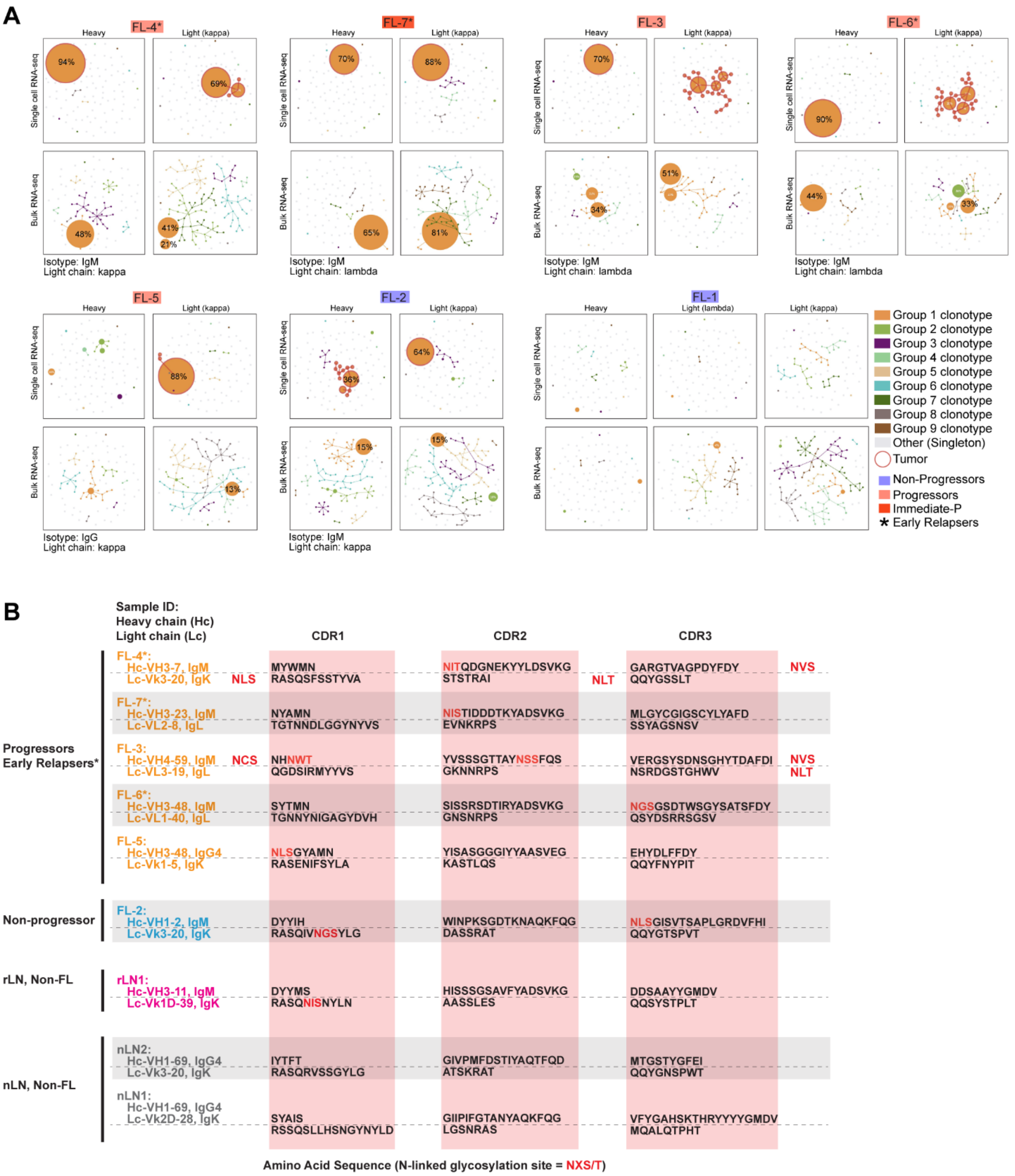


**Figure S2: RNA-seq assessment of clonotypes and *N*-glycosylation sites within variable regions of immunoglobulin genes.**

**(A)** Clonal composition maps show clonal repertoire of FL samples derived from bulk and scRNA-seq data analyses. The text below the plot indicates expressed B cell receptor as determined by bulk RNA-seq. **(B)** *N*-linked glycosylation sites in heavy and/or light chain variable regions of immunoglobulin genes from normal and malignant B cells. FL-1 is not included here due to cryptic translocation of BCL2 into the immunoglobulin locus that disrupted either expression or amplification of BCR sequences.

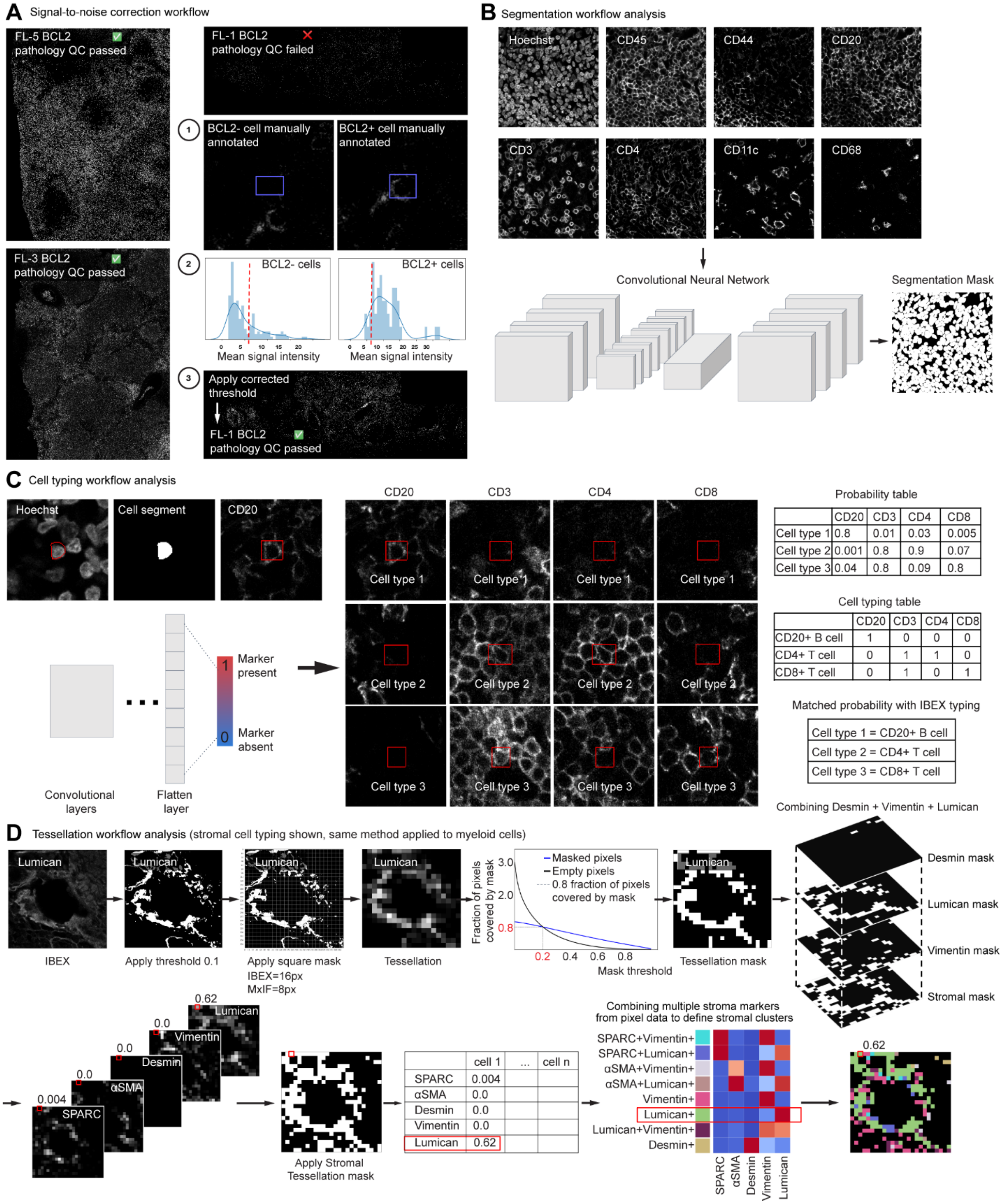


**Figure S3. Artificial intelligence (AI)-driven pipeline for multiplexed image analysis.**
**(A)** Signal-to-noise correction workflow. Representative BCL2+ images demonstrating quality control (QC) process based on signal-to-noise assessment by trained pathologists. (**1**) Example of manually annotated cells. (**2**) The distribution of mean signal intensity for BCL2- and BCL2+ cells. (**3**) Representative BCL2+ cell for patient FL-1 that now passes QC after applying the corrected threshold determined in (**2**). **(B)** Object-based cellular segmentation workflow.

Examples of the three input channels that were used: Hoechst for nuclei, CD45 for membrane, and a composite of several other markers (CD44, CD20, CD3, CD4, CD11c, CD68). Convolutional Neural Network (CNN) schema. Example of the cell segmentation mask. **(C)** Cell typing workflow analysis. Input channels: Hoechst for nuclei, cell segments, and CD20 marker, red boxes highlighting the position of the CD20+ cell. Schema of the neural network-based approach for cell typing: Convolutional layers (ResNet-50) with two neurons in the last flatten layer - marker present and marker absent. Example depicting cell type identification based on combination of markers and probability determined by CNN. In this example, four markers (CD20, CD3, CD4, CD8) from IBEX image yields three distinct cell types: Cell type 1 CD20+CD3-CD4-CD8-; Cell type 2 CD20-CD3+CD4+CD8-; Cell type 3 CD20-CD3+CD4-CD8+. CNN's output 0-1 range (probability table), where 1 corresponds to certain positive expressions and 0 corresponds to certain negative expressions (cell typing table). Matched probability with IBEX typing. **(D)** Tessellation workflow analysis (stromal cell typing shown, same method applied to myeloid cells). Original Lumican IBEX image; Lumican image after applying the threshold 0.1; Lumican image after splitting a mask derived from a marker of interest into non-intersecting squares that covers the full image area. Fraction of pixels covered by mask; Lumican image as a signal density heatmap representing the percentage of positive pixels computed for each of these squares. Tessellation masks for single markers Desmin, Lumican, Vimentin, and Stromal (Desmin+Lumican+Vimentin+) mask. An example of the markers that were used for phenotyping stromal cells from IBEX imaging data (SPARC, α-SMA, Desmin, Vimentin, Lumican). The red box highlights the square that can be roughly considered as a "pseudo-cell" measuring 16x16 pixels for stromal subpopulations identified in IBEX images. Percentages of masks in each square "pseudo-cell" is equivalent to mean cell marker expression for object-based cellular segmentation. Heatmap visualizing different stromal phenotypes. Final tessellation mask demonstrating stromal phenotypes pseudo-colored according to adjacent heatmap.

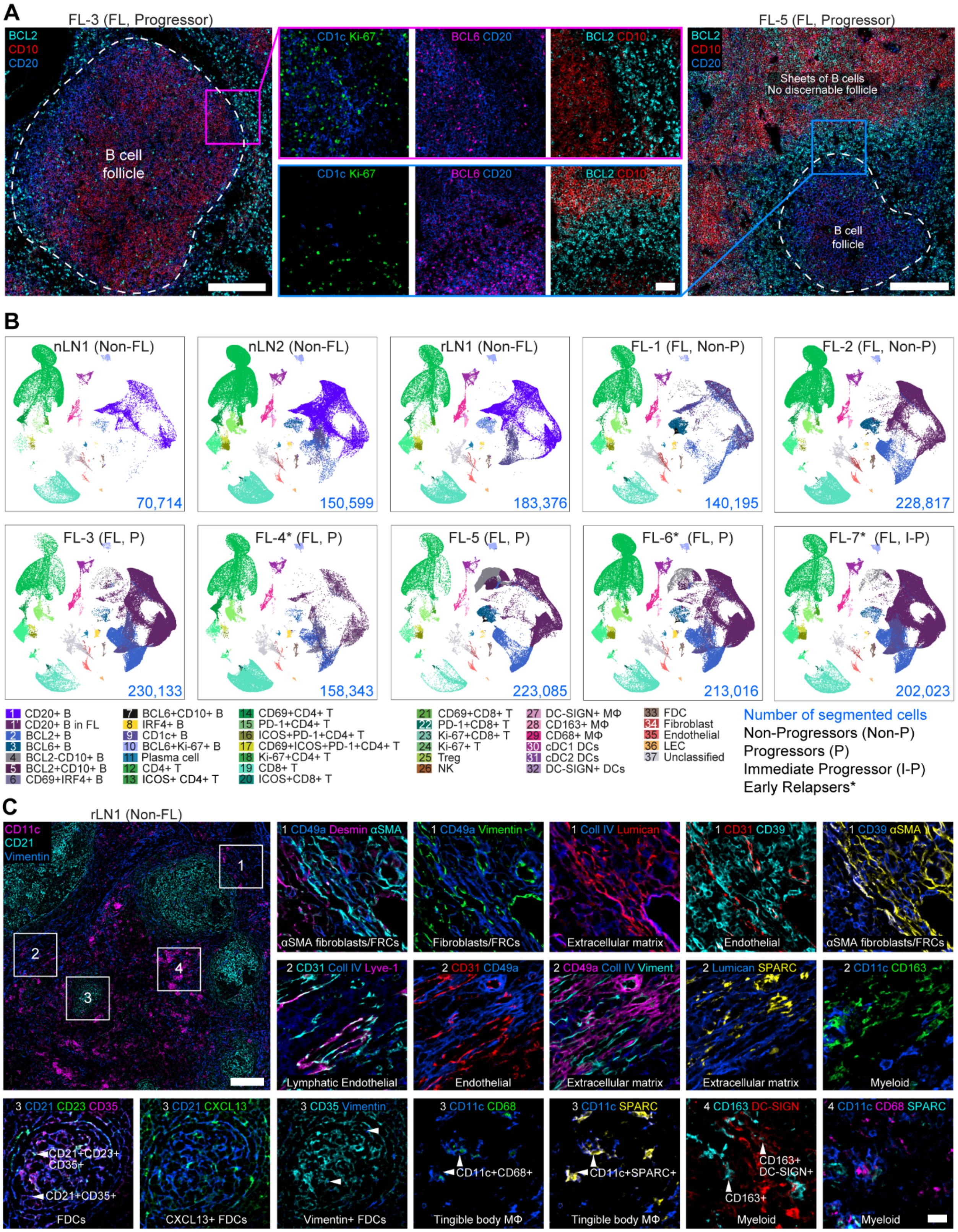


**Figure S4. IBEX imaging allows detailed spatial profiling of tumor B, myeloid, stromal, and other cell types *in situ*.**

**(A)** IBEX confocal images from two FL samples demonstrating heterogeneity in tumor B cells based on the expression of indicated markers. The selected markers are known to vary by grade and subtype of FL. Cell phenotypes based on these markers varied between samples (Figure 3D) and (**B**, see below). Scale bar is 200 μm or 50 μm (insets). **(B)** UMAP plot of single cells from individual samples colored by cell populations measured by IBEX. Cell counts per sample were obtained by object-based segmentation (light blue text). **(C)** IBEX confocal images from non-FL LN showcasing the diversity of myeloid and stromal cells present in lymphoid tissues and phenotyped using tessellation masks in Figure 3F-I. In insets, 2-3 markers are displayed per image and combined with other markers for greater clarity. Several markers are repeated to reveal different cell phenotypes. Scale bar is 200 μm or 20 μm (insets), Vimentin (Viment), Fibroblastic reticular cell (FRC), Follicular dendritic cell (FDC), single and double positive FDCs denoted by arrowheads.

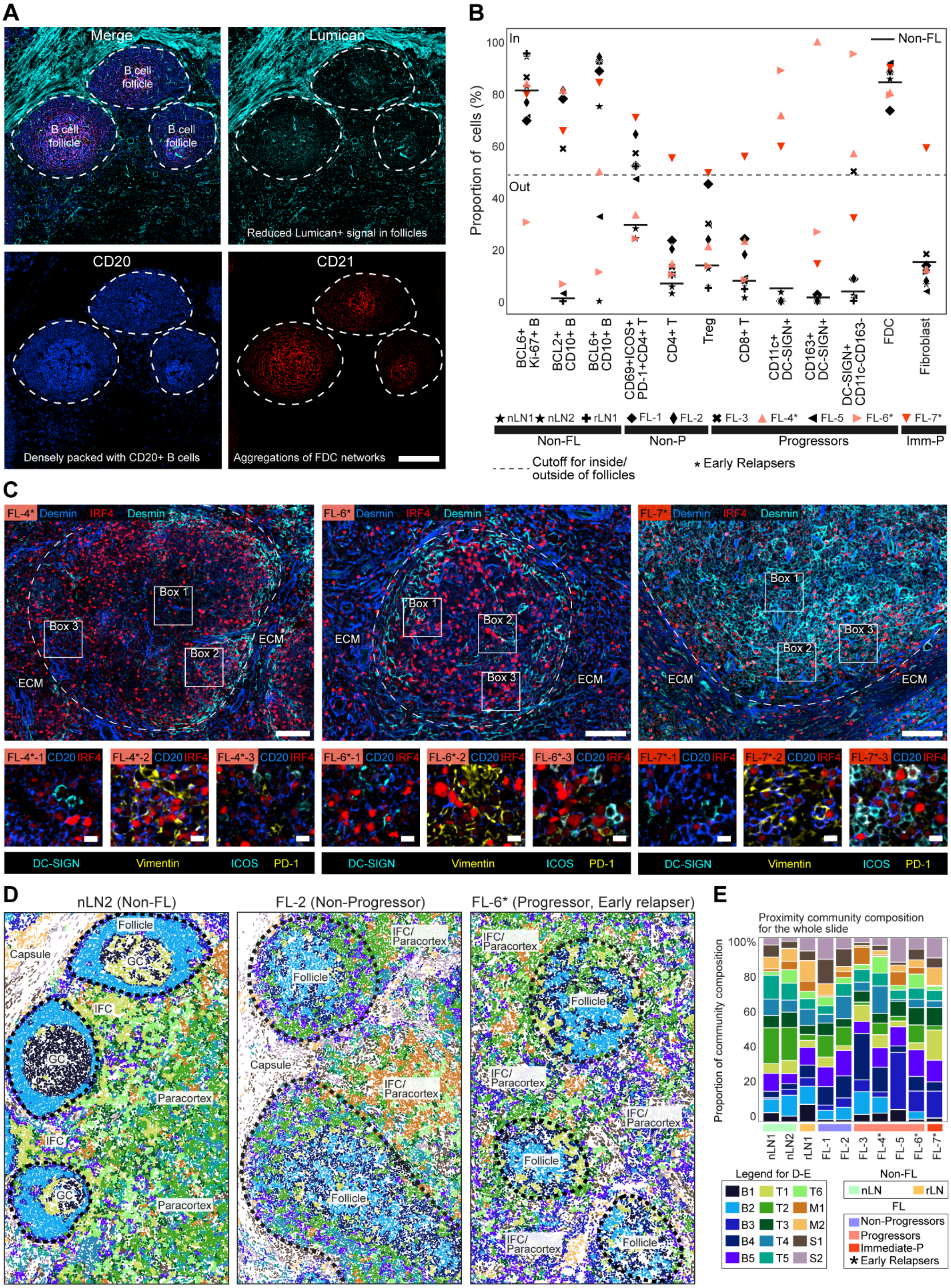
A
Merge
Lumican
B cell follicle
B cell follicle
B cell follicle
Reduced Lumican+ signal in follicles
CD20
CD21
Densely packed with CD20+ B cells
Aggregations of FDC networks
B
Non-FL
In
Out
Proportion of cells (%)
BCL6+ Ki-67+ B
BCL2+ CD10+ B
BCL6+ CD10+ B
CD69+ICOS+ PD-1+CD4+ T
CD4+ T
Treg
CD8+ T
CD11c+ DC-SIGN+
CD163+ DC-SIGN+
DC-SIGN+ CD11c-CD163-
FDC
Fibroblast
nLN1 nLN2 rLN1 FL-1 FL-2 FL-3 FL-4* FL-5 FL-6* FL-7*
Non-FL
Non-P
Progressors
Imm-P
Cutoff for inside/ outside of follicles
* Early Relapsers
C
FL-4* Desmin IRF4 Desmin
FL-6* Desmin IRF4 Desmin
FL-7* Desmin IRF4 Desmin
Box 1
Box 2
Box 3
ECM
FL-4*-1 CD20 IRF4
FL-4*-2 CD20 IRF4
FL-4*-3 CD20 IRF4
FL-6*-1 CD20 IRF4
FL-6*-2 CD20 IRF4
FL-6*-3 CD20 IRF4
FL-7*-1 CD20 IRF4
FL-7*-2 CD20 IRF4
FL-7*-3 CD20 IRF4
DC-SIGN Vimentin ICOS PD-1
D
nLN2 (Non-FL)
FL-2 (Non-Progressor)
FL-6* (Progressor, Early relapser)
Follicle
GC
Capsule
IFC
Paracortex
IFC/ Paracortex
E
Proximity community composition for the whole slide
Proportion of community composition
nLN1 nLN2 rLN1 FL-1 FL-2 FL-3 FL-4* FL-5 FL-6* FL-7*
Legend for D-E
B1 B2 B3 B4 B5
T1 T2 T3 T4 T5 T6
M1 M2 S1 S2
Non-FL
nLN rLN
FL
Non-Progressors
Progressors
Immediate-P
★ Early Relapsers

**Figure S5. Visualization and quantification of histological patterns and cellular communities using IBEX.**

**(A)** Representative IBEX images. Scale bar 200 µm. White dotted line denotes outer boundary of B cell follicles. For follicle annotations in Figures 4 and 6, B cell follicles were identified by pathologists using the presence of CD20 and CD21 and reduced density of the extracellular matrix marker lumican. **(B)** Scatter plot showing the distribution of major cell types based on IBEX image analysis. Values near zero along the y-axis represent cells located mostly outside of B cell follicles. Values near 100 represent cells located within B cell follicles. Black line indicates spatial distribution of indicated cell types for non-FL samples. **(C)** Confocal images of IRF4+ FL B cells in direct contact with DC-SIGN+ myeloid and stromal cells (Box 1), Vimentin+ FRCs (Box 2), and Tfh cells (Box 3, PD-1+ICOS+). Top row: Maximum intensity projection (MIP) of 5 µm z-stacks. Scale bar is 100 µm. Dotted line indicates B cell follicle boundary. Bottom row: Insets show 1 single z slice. Scale bar is 10 µm. **(D)** Spatial maps of selected regions of interest showing community composition of different samples pseudo-colored according to the legend. Black dotted line indicates follicle boundary. Intrafollicular cortex (IFC). The negligible IFC and paracortex in FL LNs reflects changes arising from malignancy. **(E)** Bar plots showing proportion of proximity communities identified by IBEX across the whole tissue section versus the follicles (Figure 5F).

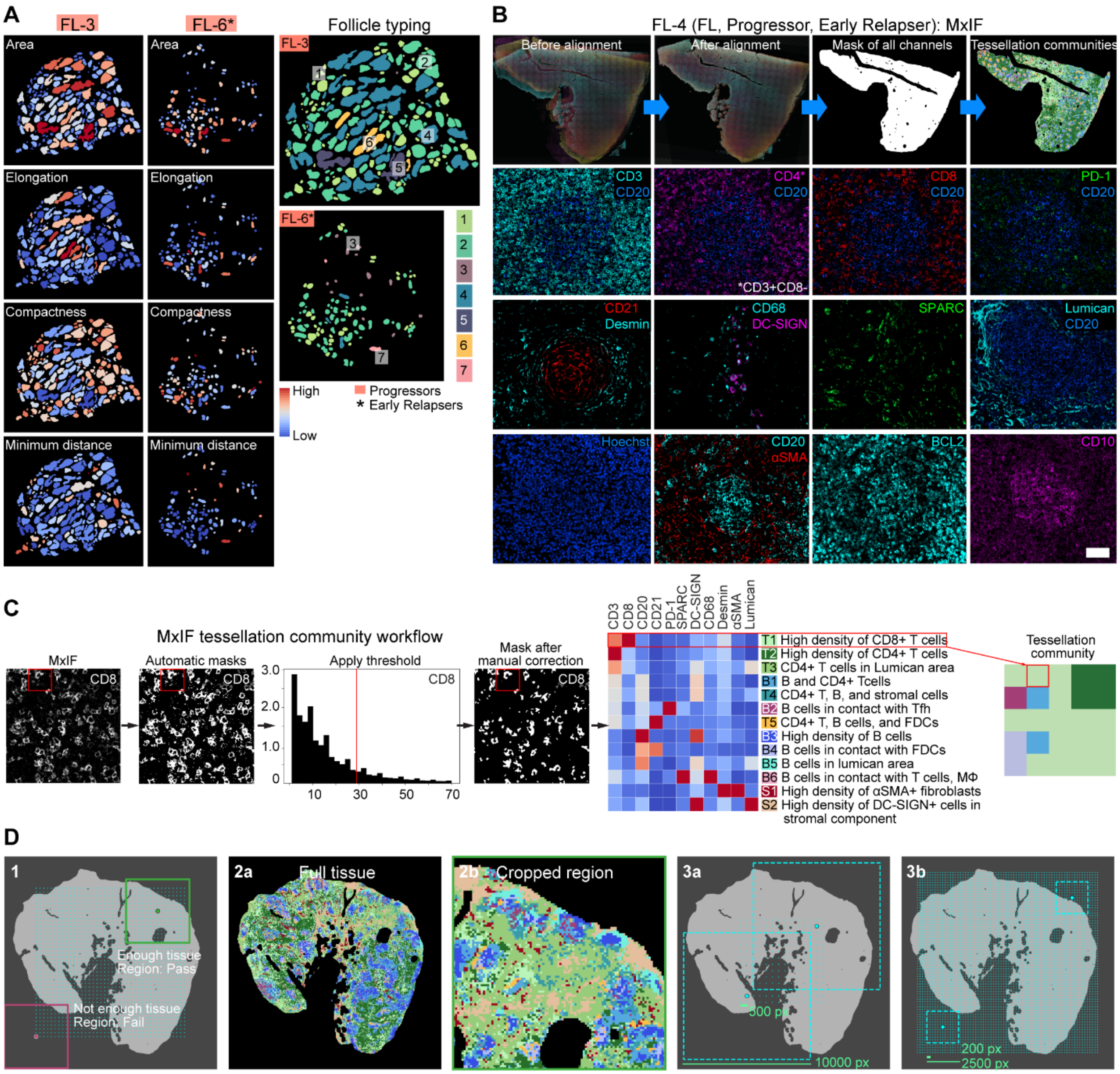


**Figure S6. Workflows for comparing spatial patterns between IBEX and MxIF images.**
**(A)** Follicle composition of MxIF images from indicated patient samples. Heatmaps depict high and low value for the size, shape, density, and distribution of B cell follicles, e.g., red follicles in area plot correspond to large follicles and dark blue correspond to small follicles. Plot showing 7 follicle types based on area, elongation, compactness, and minimum distance calculated as indicated in Figure 6C. **(B)** Top row: Workflow for serial alignment of MxIF images and assessment of tessellation communities in composite image. Confocal images of individual markers from different regions of FL-4 MxIF image. Scale bar is 50 µm. **(C)** MxIF tessellation community workflow. Original CD8 MxIF image; CD8 image after applying the automated mask; applying threshold for the image; CD8 mask after manual correction of threshold. The red box highlighted corresponds to 100 and 50 pixel-sized squares for IBEX and MxIF images respectively. Heatmap visualizing the obtained community clusters. Final tessellation mask colored as represented cellular niches. **(D)** Slide concordance analysis workflow for Figure 6. 1) Visualization of cropped images on tissue mask. The pink-boxed region contains more than 50% background and is further excluded from analysis. The green-boxed region contains more than

50% of tissue and is further compared to the full tissue area. 2) Comparison of regions including full tissue (2a) and cropped region (2b). For all regions that passed step 1, tessellation community percentages were calculated. These regions are then compared to communities present in full tissue sections using Pearson correlation. 3) Illustration of different sampling strategies based on the size of region selected. For 3a, the distance between region centers equals 500 pixels and the size of the side is equal to 10,000 pixels. For 3b the distance between region centers equals 200 pixels and the size of the side is equal to 2,500 pixels.

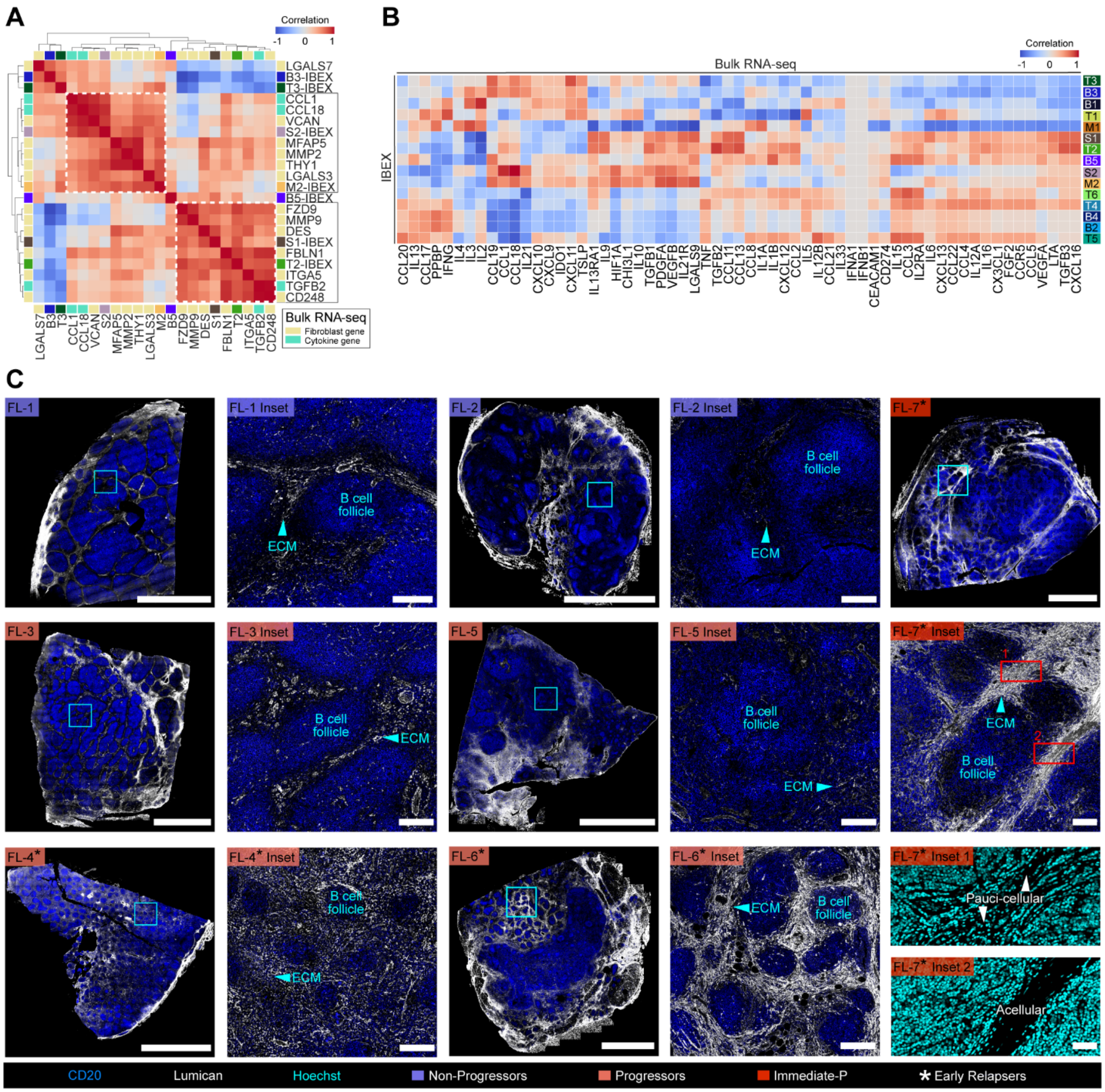


**Figure S7. Integration of multimodal technologies to evaluate normal and FL LNs.**
**(A)** Clustermap of pairwise correlations between IBEX communities (Figure 5) and gene signatures determined from bulk RNA-seq data. Colors represent the Spearman correlation coefficient. **(B)** Heatmap showing Spearman correlation between cytokine gene expression (bulk RNA-seq) and proximity community clusters (IBEX) described in Figure 5. **(C)** MxIF images showing the expansion of lumican+ extracellular matrix (ECM) around the B cell follicles of early relapsers. Scale bar is 4000 μm, 200 μm (Inset), and 50 μm (Inset 1 and 2).

### Table S1. Clinical and pathological characteristics of non-FL and FL samples.

| Patient ID | Age | Sex | BCL2 FISH | Lugano Stage | FLIPI Score | Primary Endpoint | Frontline Therapy* | After Therapy |
|---|---|---|---|---|---|---|---|---|
| nLN1 | 49 | Male | NA, non-FL | NA, non-FL | NA, non-FL | NA, non-FL | NA, non-FL | NA, non-FL |
| nLN2 | 38 | Female | NA, non-FL | NA, non-FL | NA, non-FL | NA, non-FL | NA, non-FL | NA, non-FL |
| rLN1 | 38 | Male | NA, non-FL | NA, non-FL | NA, non-FL | NA, non-FL | NA, non-FL | NA, non-FL |
| FL-1 | 31 | Male | Rearranged | 4 | 2: Intermediate risk, spontaneous remission | Non-progressor | None | NA, non-FL |
| FL-2 | 54 | Male | Rearranged | 3 | 2: Intermediate risk | Non-progressor | None | NA, non-FL |
| FL-3 | 51 | Male | Rearranged | 3 | 2: Intermediate risk | Early progressor | Lenalidomide-Rituximab | Remission |
| FL-4* | 60 | Female | Rearranged | 3 | 3: High risk | Early progressor | Bendamustine-Rituximab | Early relapser (6 months) |
| FL-5 | 71 | Male | Rearranged | 3 | 3: High risk | Early progressor | Copanlisib-Rituximab | Remission |
| FL-6* | 45 | Male | Rearranged | 4 | 2: Intermediate risk | Early progressor | Bendamustine-Rituximab | Early relapser (2 months) |
| FL-7* | 62 | Female | Rearranged | 3 | 4: High risk | Early progressor | Bendamustine-Rituximab | Early relapser (16 months) |

BCL2 Fluorescence in situ hybridization (FISH); Follicular Lymphoma International Prognostic Index (FLIPI)
*All biopsies in this study were obtained before frontline therapy.
Non-neoplastic healthy controls (nLN1, nLN2) were mesenteric LNs. The LN with reactive changes in the form of pronounced follicular hyperplasia was excised from the retroperitoneum of patient rLN1. All FL LNs, except for FL-4 (axillary) were inguinal LNs.

**Table S4. Cell-specific gene expression and cell annotations for bulk RNA-seq based deconvolution.**

| | |
|---|---|
| CD4 T cells | *ANKRD55, CD2, CD28, CD40LG, CD5, CD6, FHIT, FLT3LG, IL7R, ITK, ITM2A, KLRB1, LCK, LEF1, LRRN3, NELL2, P2RY8, TCF7, TESPA1, THEMIS, TRAT1, TRAF3IP3* |
| CD8 T cells | *CD2, CD3D, CD3G, CD3E, CD6, CD7, CD8A, CD8B, CD96, CRTAM, EOMES, FCRL6, GZMA, GZMB, GZMH, GZMK, ITK, KLRC2, KLRC4, KLRK1, PRF1, PTGDR, PVRIG, SH2D1A, TBX21, THEMIS, TIGIT, TRAC, TRAT1, TRBC2, UBASH3A, XCL2, ZAP70, CCL5, CXCR3* |
| NK cells | *CD160, CD244, CD247, CD7, CLDND2, CTSW, GZMM, IL2RB, KIR2DL1, KIR2DL2, KIR2DL3, KIR2DL4, KIR2DS2, KIR3DL1, KIR3DL2, KLRB1, KLRC2, KLRC3, KLRD1, KLRF1, KLRK1, NCAM1, NCR1, NCR3, NKG7, NMUR1, PRF1, PTGDR, PYHIN1, S1PR5, SAMD3, SH2D1B, TMIGD2, XCL2, CCL5, LIM2* |
| B cells | *BANK1, BLK, CD19, CD22, CD37, CD79A, CD79B, CLEC17A, CPNE5, CR2, FAM129C, FCRL1, FCRL2, FCRL3, FCRL5, FCRLA, HLA-DOB, MS4A1, PAX5, POU2AF1, SPIB, STAP1, TNFRSF13B, TNFRSF13C, TNFRSF17, VPREB3, CXCR5, DERL3, EAF2, FKBP11, GLCCI1, IGHG1, IGHG3, IGHM, IGKC, IGLL5, MZB1, SEC11C, SSR4, TXNDC11, TXNDC5* |
| Macrophages | *ADAP2, ADORA3, C1QA, C1QC, C3AR1, C5AR1, CCL7, CCR1, CD14, CD163, CD33, CD4, CD68, CLEC5A, CMKLR1, CSF1R, CYBB, FPR3, IL10, IL4I1, MRC1, MS4A4A, MSR1, PLA2G7, RAB7B, SIGLEC1, TREM2, VSIG4, MS4A7* |
| Monocytes | *AOAH, CCR1, CCR2, CD1D, CD300C, CD300E, CD300LB, CD302, CD33, CECR1, CSF1R, CTSS, CYBB, FCN1, IRF5, MEFV, MS4A6A, PADI4* |
| Fibroblasts | *ACTA2, ADAMTS2, CD248, COL16A1, COL1A1, COL1A2, COL3A1, COL4A1, COL5A1, COL6A1, COL6A2, COL6A3, FAP, FBLN2, FBN1, FGF2, LOXL1, MFAP5, PCOLCE, PDGFRA, PDGFRB, TAGLN, THBS2, THY1, VEGFC* |
| Endothelium | *ANGPT2, APLN, CDH5, CLEC14A, ECSCR, EMCN, ENG, ESAM, ESM1, FLT1, HHIP, KDR, MMRN1, MMRN2, NOS3, PECAM1, PTPRB, RASIP1, ROBO4, SELE, TEK, TIE1, VWF* |
| cDC | *CD1C, CD207, CLEC10A, FCER1A, HLA-DPA1, HLA-DPB1, HLA-DQB1, HLA-DRB1, HLA-DRA, ITGAX, THBD, FCGR3A, IL3RA, CLEC9A, CD14, PTPRC, ANPEP, CLEC4C, AXL, CD3G, CD19, NCAM1, SIGLEC6, CCDC103, ITGAE* |

**Table S5. Training settings for deep learning algorithms used in this study.**

| NN | Segmentation | Marker Expression | Graph Network Encoder | Graph Network Discriminator |
|---|---|---|---|---|
| Training set size | 251 images | 9,485 images | $1.8 \times 10^6$ (all cells available) | |
| Validation set size | 51 images | 1,879 images | | |
| Loss function | see (He et al., 2017) | BCE (Binary Cross Entropy) | see (Pan et al., 2018) | see (Pan et al., 2018) |
| Optimizer | SGD | Adam | Adam | Adam |
| Learning rate | 0.005 | 0.005 | 0.001 | 0.005 |
| Batch size | 4 | 15 | 1 | |
| Number of epochs | 100 | 100 | 100 | |

### Table S6. IBEX and MxIF imaging panels for fixed frozen and FFPE tissues.

| IBEX Imaging | | | | | | | |
|---|---|---|---|---|---|---|---|
| Cycle | Marker | Clone | Conjugate | Vendor | Catalog Number | Dilution | RRID |
| 1 | Hoechst | - | - | Biotium | 40046 | 1:5000 | NA |
| | CD20 | L26 | AF488 | Thermo | 53-0202-82 | 1:200 | AB_10734358 |
| | SPARC | Goat IgG | AF532 | R&D | AF941 (Unconjugated) | 1:50 | AB_2892754 |
| | CD10 | FR4D11 | PE | Caprico Biotechnologies | 103926 | 1:50 | |
| | CD10 | HI10a | PE | BioLegend | 312204 | 1:50 | AB_314915 |
| | CD3 | UCHT1 | AF594 | BioLegend | 300446 | 1:100 | AB_2563236 |
| | BCL2 | 100 | AF647 | BioLegend | 658705 | 1:25 | AB_2563279 |
| | Collagen IV<br>Goat anti-rabbit IgG | -<br>- | None<br>AF700 | Abcam<br>Thermo | Ab6586<br>A-21038 | 1:200<br>1:400 | AB_305584<br>AB_2535709 |
| 2 | Hoechst | - | - | Biotium | 40046 | 1:5000 | NA |
| | IgD | IA6-2 | AF488 | BioLegend | 348216 | 1:25 | AB_11150595 |
| | CD21 | Bu32 | AF532 | BioLegend | NA, Custom | 1:600 | AB_2892739 |
| | CD138 | MI15 | PE | BioLegend | 356504 | 1:200 | AB_2561878 |
| | CD3 | UCHT1 | AF594 | BioLegend | 300446 | 1:100 | AB_2563236 |
| | BCL6 | K112-91 | AF647 | BD Biosciences | 561525 | 1:25 | AB_10898007 |
| | CD31 | WM59 | AF700 | BioLegend | 303133 | 1:25 | AB_2566326 |
| 3 | Hoechst | - | - | Biotium | 40046 | 1:5000 | NA |
| | HLA-DR | L243 | AF488 | BioLegend | 307620 | 1:100 | AB_493175 |
| | CD23 | EBVCS-5 | AF532 | BioLegend | NA, Custom | 1:25 | AB_2892740 |
| | CD1c | L161 | PE | BioLegend | 331506 | 1:50 | AB_1088999 |
| | CD3 | UCHT1 | AF594 | BioLegend | 300446 | 1:100 | AB_2563236 |
| | CD163 | GH1/61 | AF647 | BioLegend | 333620 | 1:100 | AB_2563475 |
| | CD11c | B-Ly6 | AF700 | BD Biosciences | 561352 | 1:25 | AB_10612006 |
| 4 | Hoechst | - | - | Biotium | 40046 | 1:5000 | NA |
| | CD8 | SK1 | AF488 | BioLegend | 344716 | 1:25 | AB_10549301 |
| | CD4 | RPA-T4 | AF532 | Thermo | 58-0049-42 | 1:25 | AB_2802361 |
| | FOXP3 | 236A/E7 | eF570 | Thermo | 41-4777-82 | 1:25 | AB_2573609 |
| | CD3 | UCHT1 | AF594 | BioLegend | 300446 | 1:100 | AB_2563236 |
| | CD25 | M-A251 | AF647 | BioLegend | 356128 | 1:25 | AB_2563588 |
| | Ki-67 | B56 | AF700 | BD Biosciences | 561277 | 1:25 | AB_10611571 |
| 5 | Hoechst | - | - | Biotium | 40046 | 1:5000 | NA |
| | ICOS | CS98.4A | AF488 | BioLegend | 313514 | 1:25 | AB_2122584 |
| | CXCL13 | Goat IgG | AF532 | R&D | AF801 (Unconjugated) | 1:25 | AB_2892755 |
| | PD-1 | EH12.2H7 | PE | BioLegend | 329906 | 1:100 | AB_940483 |
| | CD3 | UCHT1 | AF594 | BioLegend | 300446 | 1:100 | AB_2563236 |
| | CD69 | FN50 | AF647 | BioLegend | 310918 | 1:25 | AB_528871 |
| 6 | Hoechst | - | - | Biotium | 40046 | 1:5000 | NA |
| | CD39 | A1 | FITC | BioLegend | 328206 | 1:50 | AB_940425 |
| | Lyve-1 | Goat IgG | AF532 | R&D | AF2089 (Unconjugated) | 1:100 | AB_2892756 |
| | CD35 | E11 | PE | BioLegend | 333406 | 1:800 | AB_2292231 |
| | CD3 | UCHT1 | AF594 | BioLegend | 300446 | 1:100 | AB_2563236 |
| | CD68 | KP1 | AF647 | Santa Cruz | sc-20060 | 1:100 | AB_2891106 |
| 7 | Hoechst | - | - | Biotium | 40046 | 1:5000 | NA |
| | α-SMA | 1A4 | AF488 | Thermo | 53-9760-82 | 1:100 | AB_2574461 |
| | Lumican | Goat IgG | AF532 | R&D | AF2846 (Unconjugated) | 1:50 | AB_2892757 |
| | IRF4 | IRF4.3E4 | PE | BioLegend | 646404 | 1:10 | AB_2563005 |
| | CD3 | UCHT1 | AF594 | BioLegend | 300446 | 1:100 | AB_2563236 |
| | DC-SIGN | 9E9A8 | AF647 | BioLegend | 330112 | 1:50 | AB_1186092 |
| 8 | Hoechst | - | - | Biotium | 40046 | 1:5000 | NA |
| | Desmin | Y66 | AF488 | Abcam | Ab185033 | 1:200 | AB_2892748 |
| | CD3 | UCHT1 | AF594 | BioLegend | 300446 | 1:100 | AB_2563236 |
| | CD49a | TS2/7 | AF647 | BioLegend | 328304 | 1:50 | AB_1236407 |
| 9 | Hoechst | - | - | Biotium | 40046 | 1:5000 | NA |
| | CD94 | DX22 | AF488 | BioLegend | 305506 | 1:50 | AB_314536 |
| | Vimentin | O91D3 | AF532 | BioLegend | Custom | 1:200 | AB_2892753 |
| | CD45 | F10-89-4 | PE/iFluor594 | Caprico Biotechnologies | 1016185 | 1:50 | AB_2892742 |
| | CD3 | UCHT1 | AF594 | BioLegend | 300446 | 1:100 | AB_2563236 |
| | CD44 | IM7 | AF647 | BioLegend | 103018 | 1:50 | AB_493681 |

| MxIF Imaging | | | | | | | | |
|---|---|---|---|---|---|---|---|---|
| Panel | Steps | Marker | Clone | Conjugate | Vendor | Catalog Number | Dilution | RRID |
| 1 | 1 | Hoechst | - | - | Biotium | 40046 | 1:5000 | NA |
| | 1 | CD20 | L26 | AF488 | Thermo | 53-0202-82 | 1:40 | AB_10734358 |
| | 1 | BCL2 | SP66 | - | Abcam | Ab236221 | 1:40 | NA |

| | | | | | | | | |
|---|---|---|---|---|---|---|---|---|
| | 2 | Donkey anti-rabbit IgG | Polyclonal | AF594 | Thermo | A-21207 | 1:200 | AB_141637 |
| | 1 | αSMA | 1A4 | eF660 | Thermo | 50-9760-82 | 1:200 | AB_2574362 |
| | 1 | CD10 | Polyclonal | - | R&D | AF1182 | 1:50 | AB_354652 |
| | 2 | Donkey anti-goat IgG | Polyclonal | AF680 | Thermo | A-21084 | 1:200 | AB_141494 |
| 2 | 1 | Hoechst | - | - | Biotium | 40046 | 1:5000 | NA |
| | 3 | Desmin | Y66 | AF488 | Abcam | Ab185033 | 1:200 | AB_2892748 |
| | 1 | CD21 | SP186 | - | Abcam | Ab240987 | 1:50 | NA |
| | 2 | Donkey anti-rabbit IgG | Polyclonal | AF555 | Thermo | A-31572 | 1:200 | AB_162543 |
| | 3 | CD68 | KP1 | iF594 | Caprico Biotechnologies | 1064135 | 1:40 | AB_2892745 |
| | 3 | DC-SIGN | h209 | - | LS Bio | LS-B3782 | 1:40 | AB_10689801 |
| | 4 | Donkey anti-rat IgG | Polyclonal | AF647 | Jackson ImmunoResearch | 712-605-153 | 1:200 | AB_2340694 |
| | 3 | SPARC | Goat IgG | - | R&D | AF941 | 1:50 | AB_355728 |
| | 4 | Donkey anti-goat IgG | Polyclonal | AF680 | Thermo | A-21084 | 1:200 | AB_141494 |
| 3 | 1 | Hoechst | - | - | Biotium | 40046 | 1:5000 | NA |
| | 5 | CD8 | EPR10640 | UT015 | Cell IDx | HI06B-005 | 1:100 | NA |
| | 6 | Anti-UT015 | Polyclonal | CL490 | Cell IDx | HI06B-005 | 1:100 | NA |
| | 3 | CD3 | SP7 | - | Abcam | Ab16669 | 1:50 | AB_443425 |
| | 4 | Goat anti-rabbit IgG | Polyclonal | AF532 | Thermo | A-11009 | 1:200 | AB_2534076 |
| | 5 | FOXP3 | 1054C | UT014 | Cell IDx | HI06B-005 | 1:100 | NA |
| | 6 | Anti-UT014 | Polyclonal | CL550 | Cell IDx | HI06B-005 | 1:100 | NA |
| | 1 | PD-1 | Polyclonal | - | Novus | AF1086 | 1:40 | AB_354588 |
| | 2 | Donkey anti-goat IgG | Polyclonal | AF680 | Thermo | A-21084 | 1:200 | AB_141494 |
| 4 | 1 | Hoechst | - | - | Biotium | 40046 | 1:5000 | NA |
| | 1 | Lumican | Goat IgG | AF532 | R&D | AF2846 (Unconjugated) | 1:50 | AB_2892757 |
| | 2 | Donkey anti-goat IgG | Polyclonal | AF555 | Thermo | A-21432 | 1:200 | AB_2535853 |

The following markers were excluded from tessellation-based analysis due to poor antibody performance and/or high background: BCL6, CD4, CD20 (in Panels 2-4), CD34, HLA-DR, IRF4, Ki-67, and Vimentin.

**Table S7. Bulk RNA-seq gene signatures used for cell community assessment (Figure 7C-D, Figure S7B-C)**

| **Cytokine signature name** | **Cytokine gene list** |
|---|---|
| Matrix metalloproteinases | *MMP2, MMP3* |
| (Myo)fibroblasts | *FAP, CD248, ACTA2* |
| Matrix collagen | *COL11A1, COL4A1, COL5A1, COL1A2, COL3A1, COL1A1* |
| Mesenchymal cells | *NT5E, THY1, ITGA5, VCAM1, NOTCH3, FZD9, ENG* |
| Cancer cell migration | *LAMA3, LAMC2, LAMB3* |
| Fibroblastic reticular cells | *NT5E, LTBR, ICAM1, DES, ACTA2, PDPN, PTGS2, VIM, VCAM1, PDGFRA, THY1* |
| T cell exhaustion | *TIGIT, PDCD1LG2, PDCD1, HAVCR2, BTLA, CTLA4, LAG3, CD274* |
| B cell antibody production | *IFNB1, IFNG, IL2, IL21, IL21R, IL15, IL6, IL10* |
| Th2 response | *IL4, IL6, IL10, CCL8, CXCL13* |
| Myeloid inflammation | *IL1A, IL1B, TNF, IL6, LTA* |
| Treg recruitment | *CXCL13, CCL1, CCL3, CCL4, CCL5, CCL18, CCL17, CCL22, CCL19, CCL21* |
| Th1 response | *IFNG, IFNA1, IFNB1, IL12A, IL12B, IL2, TNF, IL9* |
| Granulocyte activation | *IL5, IL9, CCL13* |
| NK cell recruitment | *IL15, CCL5, CCL3, CCL4, CXCL9, CXCL10, CCL19, CCL21* |
| T cell activation | *TGFB1, TGFB3, IL12, CXCL9, IFNG, LGALS9, CEACAM1, CD274* |
| Th1/CD8+ T cell recruitment | *CCL3, CCL4, CXCL9, CXCL10, CCL5, CCL19, CCL21* |
| B cell recruitment | *IL12A, IL12B, CCL19, CCL20, CCL21, CXCL13, CXCL12, CXCL10* |
| Fibroblast recruitment | *CCL4, CCL2, FGF2, PDGFA, TGFB1, TGFB3, HIF1A, VEGFA, VEGFB, CXCL12, IL6* |

Gene signatures were manually annotated from several sources (Jonigk et al., 2019; Kamp et al., 2022; Luzina et al., 2015; Mourcin et al., 2021; Nagarsheth et al., 2017).
